# Functional and ultrastructural analysis of reafferent mechanosensation in larval zebrafish

**DOI:** 10.1101/2021.05.04.442674

**Authors:** Iris Odstrcil, Mariela D. Petkova, Martin Haesemeyer, Jonathan Boulanger-Weill, Maxim Nikitchenko, James A. Gagnon, Pablo Oteiza, Richard Schalek, Adi Peleg, Ruben Portugues, Jeff Lichtman, Florian Engert

**Affiliations:** Department of Molecular and Cellular Biology, Faculty of Arts and Sciences, Harvard University, Cambridge, Massachusetts, 02138, USA; Center for Brain Science, Faculty of Arts and Sciences, Harvard University, Cambridge, Massachusetts, 02138, USA; The Ohio State University, Department of Neuroscience, Columbus, Ohio, 43210, USA; Duke University School of Medicine, Durham, North Carolina, 27707, USA; School of Biological Sciences, University of Utah, Salt Lake City, Utah, 84112, USA; Center for Cell & Genome Science, University of Utah, Salt Lake City, Utah, 84112, USA; Max Planck Institute for Ornithology, Flow Sensing Research Group, Seewiesen, 82319, Germany; Institute of Neuroscience, Technical University of Munich, Munich, 80333, Germany; Max Planck Institute of Neurobiology, Research Group of Sensorimotor Control, Martinsried, 82152, Germany; Munich Cluster for Systems Neurology (SyNergy), Munich, 81377, Germany; Friedrich Miescher Institute for Biomedical Research, Basel, 4056, Switzerland

## Abstract

All animals need to differentiate between exafferent stimuli, which are caused by the environment, and reafferent stimuli, which are caused by their own movement. In the case of mechanosensation in aquatic animals, the exafferent inputs are water vibrations in the animal’s proximity, which need to be distinguished from the reafferent inputs arising from fluid drag due to locomotion. Both of these inputs are detected by the lateral line, a collection of mechanosensory organs distributed along the surface of the body.

In this study, we characterize in detail how the hair cells, which are the receptor cells of the lateral line, discriminate between such reafferent and exafferent signals in zebrafish larvae. Using dye labeling of the lateral line nerve, we visualize two parallel descending inputs that can influence lateral line sensitivity. We combine functional imaging with ultra-structural EM circuit reconstruction to show that cholinergic signals originating from the hindbrain transmit efference copies that cancel out self-generated reafferent stimulation during locomotion, and that dopaminergic signals from the hypothalamus may have a role in threshold modulation both in response to locomotion and salient stimuli. We further gain direct mechanistic insight into the core components of this circuit by loss-of-function perturbations using targeted ablations and gene knockouts.

We propose that this simple circuit is the core implementation of mechanosensory reafferent suppression in these young animals and that it might form the first instantiation of state-dependent modulation found at later stages in development.

## Introduction

To respond appropriately to their environment, animals must distinguish between reafferent signals generated by their own motion and exafferent sensory inputs caused by external sources. To avoid this informational ambiguity, sensory pathways may be completely blocked during locomotion, or reafferent inputs may be precisely subtracted out to allow for the isolation and transmission of exafferent signals. An early observation of this phenomenon was made by Hermann von Helmholtz, who noted that our ability to process visual images is severely compromised during saccadic eye movements and suggested that visual motion information is transiently suppressed during these motor events (von Helmholtz, 1867). Since then, examples of both complete sensory suppression and reafferent cancellation have been found for multiple modalities and across species throughout the animal kingdom (see Crapse and Sommer, 2008; Cullen, 2004; Straka et al., 2018 for review). Examples include the suppression of reafferent sounds in zebra finches and mice (Keller and Hahnloser, 2009; Schneider et al., 2014, 2018), suppression of reafferent somatosensation during crayfish escapes or fish respiratory and fin movements (Bryan and Krasne, 1977; Perks et al., 2020); and the subtraction of visual flow during body saccades in *Drosophila* (Kim et al., 2015). At their core, all these implementations rely on corollary discharge, or efference copy, signals from motor-command centers that inform the sensory processing pathway about impending movements (von Holst and Mittelstaedt, 1950; Sperry, 1950).

One of the earliest detailed dissections of the neural mechanisms underlying reafferent cancellation was accomplished in weakly electric fish, so named for their ability to emit electrical discharges. Electrosensory receptor organs located in the lateral line detect the resulting electric field lines, and distortions in these fields are used for electrolocation by dedicated downstream circuits in the brain. Notably, the self-generated electrical pulses themselves are adaptively cancelled out by cerebellar-like structures precisely tuned to subtract a copy of the expected incoming sensory information in a flexible and dynamic fashion (Bell, 1981; Montgomery and Bodznick, 1994; Zipser and Bennett, 1976).

Most fish and amphibia lack such specialized electrosensory organs and use the lateral line exclusively to sense water motion relative to their bodies (Dijkgraaf, 1963). This information can be used to identify moving animals in their vicinity, as well as abiotic water currents, and therefore contributes to a variety of behaviors including schooling (Pitcher et al., 1976), prey capture (Montgomery et al., 2002), predator avoidance (Stewart and McHenry, 2010), and rheotaxis (Oteiza et al., 2017; Suli et al., 2012). Because fluid drag during locomotion strongly activates the lateral line, in this case too, external and self-generated stimuli need to be processed differentially.

Indeed, the transient inhibition of the afferent lateral nerve during locomotor events has been observed in tadpoles (Chagnaud et al., 2015; Russell, 1971), and diverse fish species (Russell, 1976; Russell and Roberts, 1974), including zebrafish (Lunsford et al., 2019; Pichler and Lagnado, 2020). While the general characteristics of this phenomenon appear to be maintained across aquatic animals, the specific details regarding precise connectivity, anatomy and function vary depending on the organism under inquiry (see Coombs et al., 2012) for a comprehensive review).

Specifically in the zebrafish, two efferent pathways that target the lateral line have been identified. One originating from the cholinergic Octavolateral Efferent Nucleus (OEN) in the hindbrain and the other from the Dopaminergic Efferent to the Lateral Line (DELL) in the ventral hypothalamus (Bricaud et al., 2001; Metcalfe et al., 1985). Further, acetylcholine has been shown to inhibit hair cell activity (Carpaneto Freixas et al., 2021; Dawkins et al., 2005), making the OEN the most likely source of reafferent suppression. At the synaptic level, ideas about how acetylcholine may inhibit hair cells come from work done in the mammalian inner ear. In the rat cochlea, the hyperpolarizing effects of acetylcholine result from the activation of nicotinic receptors containing specialized α9/α10 subunits (Elgoyhen et al., 1994, 2001). This inhibitory effect, however, is less related to reafferent suppression and is thought to have a protective function when animals are exposed to acoustic hyperstimulation that can induce damage due to excitotoxicity (Schneider et al., 2014, 2018; Taranda et al., 2009). Interestingly, enriched expression of the α9, but not the α10 subunits, have been observed in the hair cells of the lateral line of larval zebrafish (Carpaneto Freixas et al., 2021; Erickson and Nicolson, 2015), but whether they play a specific role in protective silencing, context-dependent modulation or treafferent suppression has not yet been determined. Nonetheless, there is now significant evidence for the role of cholinergic efferents in reafferent suppression, and their mechanism has been hypothesized to the level of individual receptor subtypes. The functional role and mechanism of the dopaminergic efferent neurons, on the other hand, is still very much unclear.

It is further unknown how the convergence of all of the involved cell populations, namely the two efferent subtypes (OEN and DELL), the afferent sensory neurons, and the hair cells of the neuromasts, gives rise to an interconnected and complex microcircuit that can adaptively process the sensory signals detected by the lateral line.

In this study, we address these questions by exploiting the small size and optical accessibility of the larval zebrafish, which afford a unique opportunity to perform an exhaustive anatomical and functional dissection of the complex microcircuitry surrounding the neuromasts of the lateral line. To that end, we first described the identity and morphology of descending inputs to the lateral line. Further, we use a combination of confocal and electron microscopy to unveil the detailed synaptic structures between efferent terminals, afferent neurites and the hair cells of the neuromast. Using functional imaging of all neuronal populations during behavior, we observe activation of both OEN and DELL neurons during locomotion, with a concomitant and significant reduction in afferent activity. However, targeted laser ablation of individual populations, established that OEN, but not DELL, neurons are necessary for the suppression of reafferent activity. We further confirm the critical role of the cholinergic pathway and establish the necessity of the α9 cholinergic subunit in hair cells by means of gene knockout approaches. Additionally, we observed that DELL, but not OEN neurons, respond to flow, acoustic, and visual stimuli in the absence of motor outputs—unveiling a sensory capacity in these dopaminergic neurons that could be used to modulate mechanosensation in response to environmental cues.

Taken together our results identify the OEN as the source of efference copy signaling to the lateral line, and they allow us to propose the hypothesis that DELL neurons serve to modulate hair cell sensitivity during the quiescent periods following locomotion and after the occurrence of salient stimuli.

## Results

### Anatomy

If efference copy mechanisms are in place to cancel out self-generated mechanical stimulation during locomotion, there must exist an anatomical bridge between the motor pathway and the sensory stream of the lateral line. This connection could exist at any level within both the motor and sensory pathways (Figure 1A). In the larval zebrafish, descending axons from the brain have been found to directly innervate the peripheral mechanosensory organs, the neuromasts (Metcalfe et al., 1985). These axons reach the tail neuromasts via the posterior lateral line nerve, a superficial nerve that extends along the myoseptum and comprises the dendrites of primary sensory neurons (Figure 1B). To visualize the neurons that make up the lateral line nerve, we injected fluorescent dextrans at a rostral position within the nerve. Consistent with previous studies, this procedure successfully labeled both the afferent and the efferent neuronal populations that target the neuromasts of the lateral line: the primary sensory neurons of the posterior lateral line ganglion (PLLg), and the efferent cells, which are found in two small clusters: the Diencephalic Efferent to the Lateral Line (DELL) in the hypothalamus and the Octavolateral Efferent Nucleus (OEN) in the hindbrain (Figures 1C and 1D) (Bricaud et al., 2001; Metcalfe et al., 1985).

**Figure 1.**
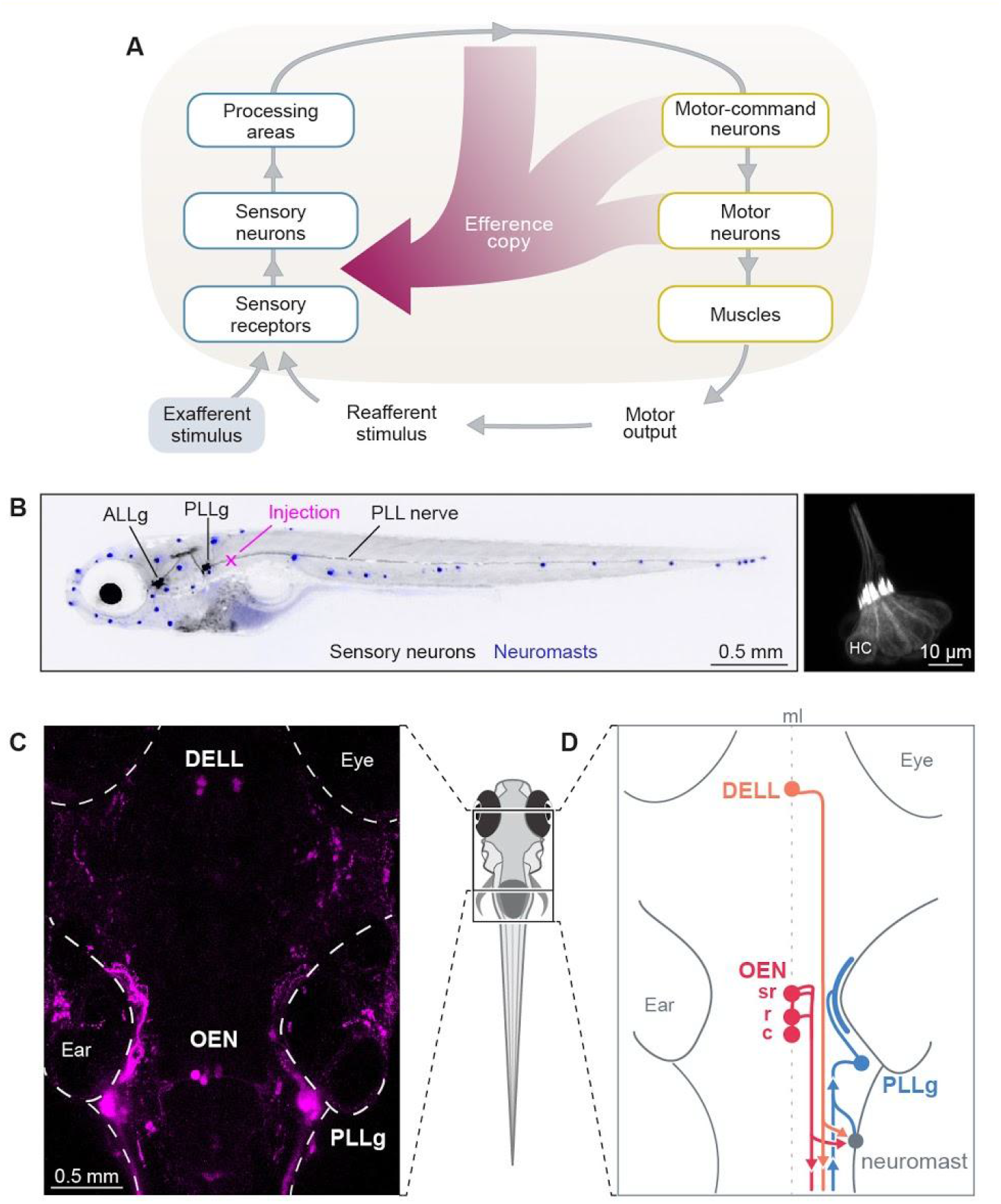
Efference copy signals and lateral line circuitry. **(A)** Schematic of a sensorimotor circuit comprising a sensory (blue) and motor pathway (yellow). Each pathway is organized hierarchically, where higher tiers denote increased processing complexity and distance from the periphery. Across the animal kingdom, efference copy signals (crimson) or other types of corollary discharges can arise from almost all levels of the motor pathway and target any level in the sensory pathway. In the larval zebrafish, centrifugal fibers from the brain project directly to the neuromasts (sensory receptors/ neurons). Adapted from Crapse and Sommer, 2008. **(B)** Left: lateral view of a 7 dpf Hgn39D:GFP zebrafish larva expressing GFP in lateral line primary neurons (black). Hair cells have been stained using the vital dye DiASP, revealing neuromasts as blue dots. Backfills of the lateral line nerve were performed by injecting dyes at the level of the second myotome (magenta cross). Right: close-up of a posterior neuromast expressing GFP in its component hair cells (HC) of a 7 dpf larva. **(C)** Maximum intensity projection of the brain of a 6 dpf larva that received bilateral injections of fluorescent dextrans in the PLL nerve. The labeled neurons cluster into 3 main cell nuclei: the PLL ganglion (PLLg), the Diencephalic Effrent of the Lateral Line (DELL), and the Octavomedial Efferent Nucleus (OEN) with its caudal, rostral and supra-rostral subdivisions. **(D)** Diagram showing the major components of the PLL circuit. Neuromasts, peripheral sensory organs (gray), are activated by mechanosensory stimuli. Upon neuromast activation, signals are transmitted to primary sensory neurons/afferents whose cell bodies cluster in the PLLg (light-blue) and whose axonal projections extend to the hindbrain. Descending inputs originate from the DELL in the ventral hypothalamus (orange) and from the OEN in the hindbrain (red). The circuit is midline (ml) symmetric but only one side is illustrated for clarity.

The OEN has been further subdivided into rostral and caudal subnuclei, according to the relative positions of the soma of the cells that comprise them, and the location at which their axons exit the brain (Figures 1D and S1A) (Metcalfe et al., 1985). Dye labeling of multiple animals and subsequent registration of the images onto a reference brain, confirmed these anatomical subdivisions and revealed that OEN neurons also occupy a third position in the hindbrain at the level of rhombomere 4 (Figure S1A), which we have termed the supra-rostral OEN (srOEN).

To further characterize the system, we sought to confirm the primary neurotransmitter identity of both nuclei by performing dextran injections in transgenic fish lines that label dopaminergic (DAT:GFP), mono-aminergic (ETvmat2:GFP) or largely cholinergic (Isl1:GFP) neurons. Since Isl1 is expressed by a variety of neurons, we supplemented these injections with immunohistochemical stains against choline acetyltransferase. In line with previous studies, we observed that DELL neurons are dopaminergic and OEN neurons are cholinergic (Figures S1B and S1C) (Haehnel-Taguchi et al., 2018; Xi et al., 2011). Furthermore, these experiments allowed us to ascertain that the above-mentioned transgenic lines label the efferent neurons of the lateral line, and validated them as anatomical markers for subsequent experiments.

To map the patterns of efferent innervation and understand the organization principles of this circuit, we performed single-cell focal electroporations to label individual neurons with membrane-tagged fluorescent proteins, and analyzed their arborization patterns. We focused on describing neuronal morphology and further used this description to ask whether efferent nuclei are somatotopically organized, that is, whether the position of a neuron’s soma within the nucleus correlates with the rostro-caudal distribution of its recipient neuromasts. We found that neither DELL nor OEN appear to be strictly somatotopically organized (Figure S1E). Individual DELL neurons extend neurites ipsilaterally and innervate a wide range of targets: neuromasts of the head and tail, hair cells of the inner ear, as well as the spinal cord (Figure S1D, top, and S1E) (Jay et al., 2015; McPherson et al., 2016; Tay et al., 2011). Individual OEN neurons, by contrast, possess bilateral dendritic arbors. Their axons, however, project ipsilaterally, and contact hair cells of the inner ear, and of anterior and posterior neuromasts (Figures S1D and S1E). Any combination of targets other than the inner ear alone have been observed (Figure S1E). Notably, neurons belonging to the different OEN subnuclei have overlapping targets, indicating that neuromast innervation is not governed by the efferent’s soma position in the hindbrain (Figures S1D, middle and bottom, and S1E).

Overall, tracing of individual efferent neurons revealed a large degree of divergence that is not somatotopically organized. Since neuromasts in different parts of the body can receive inputs from the same efferent neurons and will likely experience temporally synchronized stimuli during swimming, it is likely that the efferent mechanisms in place act in bulk, rather than being finely tuned to different regions of the body.

In summary, we confirmed the existence of two sources of descending inputs to the hair cells of the lateral line of zebrafish larvae. The first, the DELL, is a dopaminergic hypothalamic nucleus and the second, the OEN, is a cholinergic nucleus with further anatomical subdivisions. Both nuclei are anatomically well-poised to provide sensory modulation, and could transmit efference copy signals to compensate for self-generated stimulation during locomotion.

### Functional properties of afferent and efferent neurons during locomotion

#### Afferent neurons in the posterior lateral line ganglion respond to exafferent mechanosensory stimuli but not to reafferent stimulation

If efference copies act directly on the sensory pathway rather than at later processing stages, then their effects should be seen in the output of the primary sensory neurons in the PLLg. To test this hypothesis, we combined stimulus delivery and behavioral tracking with 2-photon functional imaging of PLLg neurons in transgenic Tg(elavl3:GCaMP6s/f) zebrafish larvae expressing fluorescent calcium indicators under the control of a neuronal promoter. In our assay, larvae were restrained in an agarose polymer but were free to move their tail (Figure 2A). We delivered brief local flow currents by water injections through a micropipette and observed that, as expected, sensory neurons are responsive to external flow stimuli (Figures 2B-D) (Liao, 2010). It has been previously observed that most hair cells of the larval posterior lateral line are directionally tuned to two flow directions: head-to-tail or tail-to-head (Ghysen and Dambly-Chaudière, 2007; López-Schier et al., 2004; Pichler and Lagnado, 2019), and that afferent sensory neurons exclusively innervate similarly-tuned hair cells (Dow et al., 2015; Faucherre et al., 2009; Nagiel et al., 2008). Consistent with this anatomical architecture, we found that head-to-tail flow activated only a subset of sensory neurons in the PLLg, presumably the fraction that is specifically tuned to that direction (Figure 2C). It is worth noting that we used head-to-tail stimulation, because this is the main direction of swim-induced flow since zebrafish larvae rarely swim backwards.

**Figure 2.**
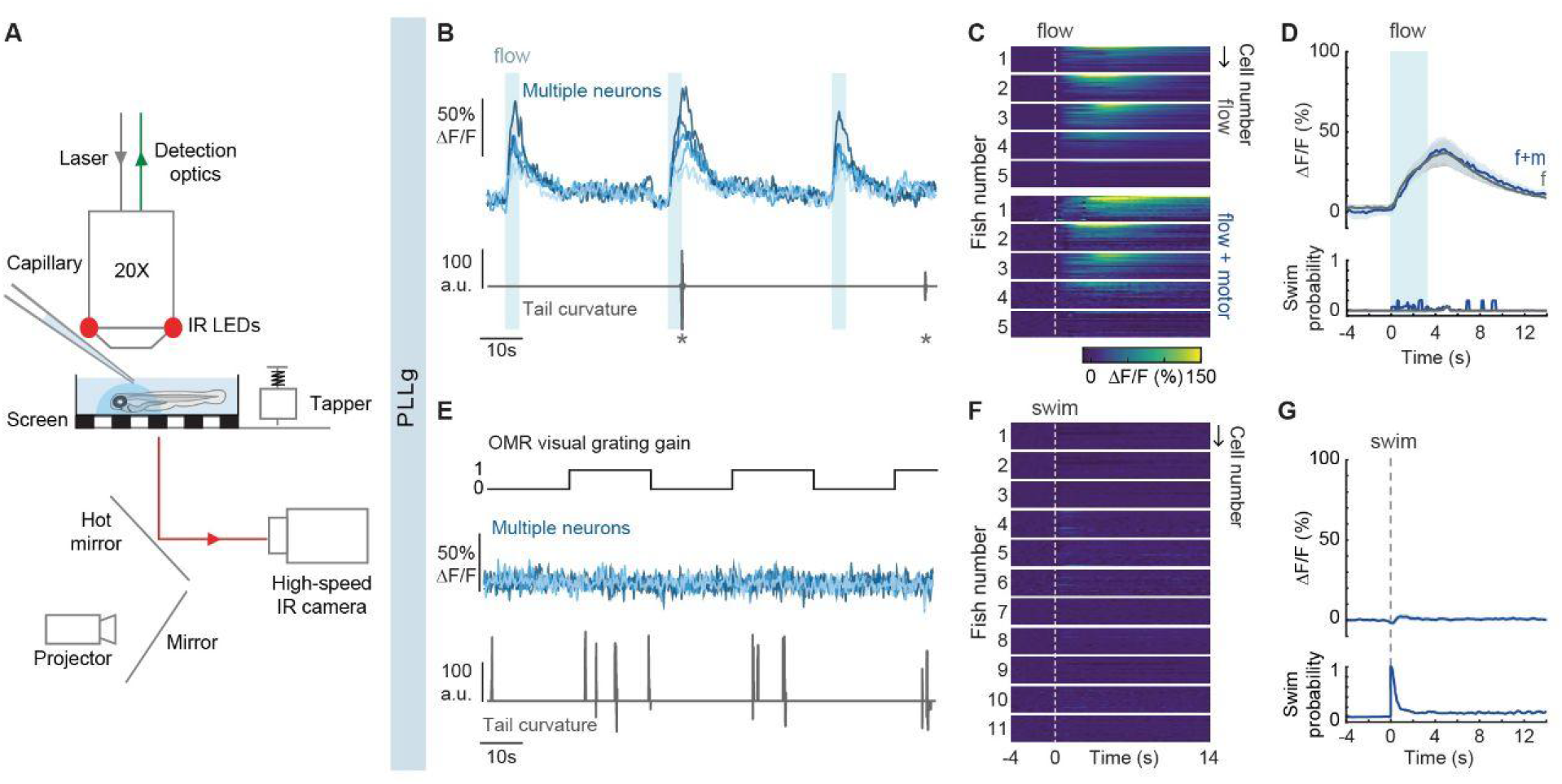
Functional imaging of primary sensory neurons in the PLLg during head-restrained swimming. **(A)** Illustration of the experimental setup. Zebrafish larvae are embedded in agarose but are able to move their tail. 2-photon imaging is performed to record calcium-dependent fluorescence signals while tail position is simultaneously tracked using an infrared (IR) camera. Visual stimuli are projected onto a diffusive screen under the animal, taps result from the impact of an electronically controlled solenoid against the stage, and flow is delivered by pressure injection through a micropipette positioned above the animal’s head. **(B)** Example traces of top: fluorescence activity (ΔF/F) of 5 responsive neurons in the PLLg of a single fish expressing GcAMP6s during flow delivery (light blue), and bottom: the animal’s concomitant cumulative tail curvature. Swim bouts are marked with stars. **(C)** Average single-cell responses to 3.3 s of water flow delivery separated into instances that did and did not elicit motor responses. (Flow: n= 24, 30, 20, 26, 16 cells in 5 fish, Flow + motor: n= 23, 15, 18, 15, 16 cells of the same fish). Flow onset indicated by the dotted line. Since stimulus strength was calibrated to trigger behavior stochastically, some neurons were only imaged during flow delivery instances that did not result in motor output, hence the discrepancy in cell number within individual fish across conditions. **(D)** Population average of stimulus-triggered neuronal activity in the PLLg (mean ΔF/F ± s.e.m.) in response to water flow that elicited (f+m, blue) and did not elicit (f, gray) motor responses (n= 5 fish). Averages arise from the single-cell measurements shown in Figure 2C. While the flow delivery period lasted 3.3 s, it is possible that a residual stimulus persisted after flow delivery was terminated. Bottom: swim probability during the same time period. **(E)** Example traces of top: fluorescence activity (ΔF/F) of 5 representative neurons in the PLLg of a single fish expressing GcAMP6s. A black and white grating was projected either statically or moving caudo-rostrally with respect to the fish at alternating 20 s intervals to promote swimming. Bottom: the animal’s concomitant cumulative tail curvature. **(F)** Average single-cell responses (ΔF/F) during swim bouts in 11 fish. (n= 22, 16, 19, 13, 17, 23, 27, 25, 22, 26, 23 cells). Swims start at dotted line. **(G)** Population swim-triggered averages of neuronal activity in the PLLg (mean ΔF/F ± s.e.m.). Bottom: swim probability during the same period. Averages arise from the single-cell measurements shown in Figure 2F.

In order to dissect the relative contributions of reafferent and exafferent flow stimuli to sensory neuron activity, we adjusted the flow strength to evoke swim events sporadically, enabling us to separate trials into swim and no-swim categories. In cases where swim events occurred during flow delivery, hair cells were stimulated by two separate sources: flow and locomotion. Nonetheless, sensory neuron activity was indistinguishable from that induced by exafferent flow alone (Figures 2C and 2D). This suggests that the reafferent flow stimulation is balanced by efferent inhibition, and that concurrent exafferent flow stimuli can evoke additional activity.

To complement this observation, we measured the responses of lateral line sensory neurons during swimming in the absence of exafferent stimulation. Since zebrafish larvae have a low motor drive in our head-embedded preparation, we used a black and white moving grating to elicit the optomotor response (OMR) (Neuhauss et al., 1999; Orger et al., 2008). In this way, we focused our analysis on the sensory activity during visually-evoked swim events where the flow patterns detected by the lateral line are exclusively generated by the fish’s own motion (reafferent stimulation) (Figure 2E). Remarkably, even though hair cells are strongly deflected by fluid drag during tail undulations (Cahn and Shaw, 1965), we observed that self-induced flow stimuli did not result in sensory neuron activation during locomotion (Figures 2F and 2G).

These experiments show that whilst sensory neurons in the PLLg can be activated by exafferent mechanosensory stimuli, they are not excited by hair cell deflections during swims, favoring the hypothesis that efference copy signals exist in the lateral line system of larval zebrafish, and that these are transmitted directly to the peripheral sensory pathway.

#### DELL and OEN neurons exhibit graded motor-correlated activity, while DELL neurons are also activated by sensory stimuli

A hallmark of efference copy signals is their temporal coincidence with motor outputs. We asked whether DELL or OEN neurons exhibit this feature by monitoring their activity using the functional imaging assays described above. In addition to OMR-induced swimming, we extended our survey of locomotor outputs to include startle responses. A long-standing hypothesis in the field has proposed that since swims and escapes are governed by distinct motor-command centers (Orger et al., 2008; Ritter et al., 2001; Severi et al., 2014), they could in theory, recruit dedicated efferent populations differentially (Kimmel et al., 1974). To address this directly, we evoked startle responses with a mechanical tapper that struck the microscope’s stage and whose strength was again calibrated to evoke motor responses sporadically.

We found that activity in hypothalamic DELL neurons was locked to the onset of locomotion both during swims and startle responses (Figures 3A-B, 3D and S2A-B). Interestingly, DELL neurons were also activated, albeit less strongly, by visual stimuli and taps that did not evoke motor outputs, unmasking a purely sensory component to their activity (Figures 3C, 3D and S2B) (Reinig et al., 2017). In line with these results, swims evoked by visual stimulation elicit a stronger response than spontaneous swim events (Figure S2C). To further probe the sensitivity of DELL neurons to other sensory modalities, we delivered flow and heat stimuli while continuing to synchronously monitor motor and neural function. DELL neurons strongly responded to flow but not to heat, indicating that this nucleus cannot be recruited by all sensory modalities (Figures 3E and 3F). Furthermore, we found that DELL sensitivity to mechanical stimuli does not only originate from the lateral line, but also depends on the inner ear, since activation by taps is preserved after specifically ablating the hair cells of the lateral line by external copper-sulfate application (Figure S2D) (Lacoste et al., 2015).

**Figure 3.**
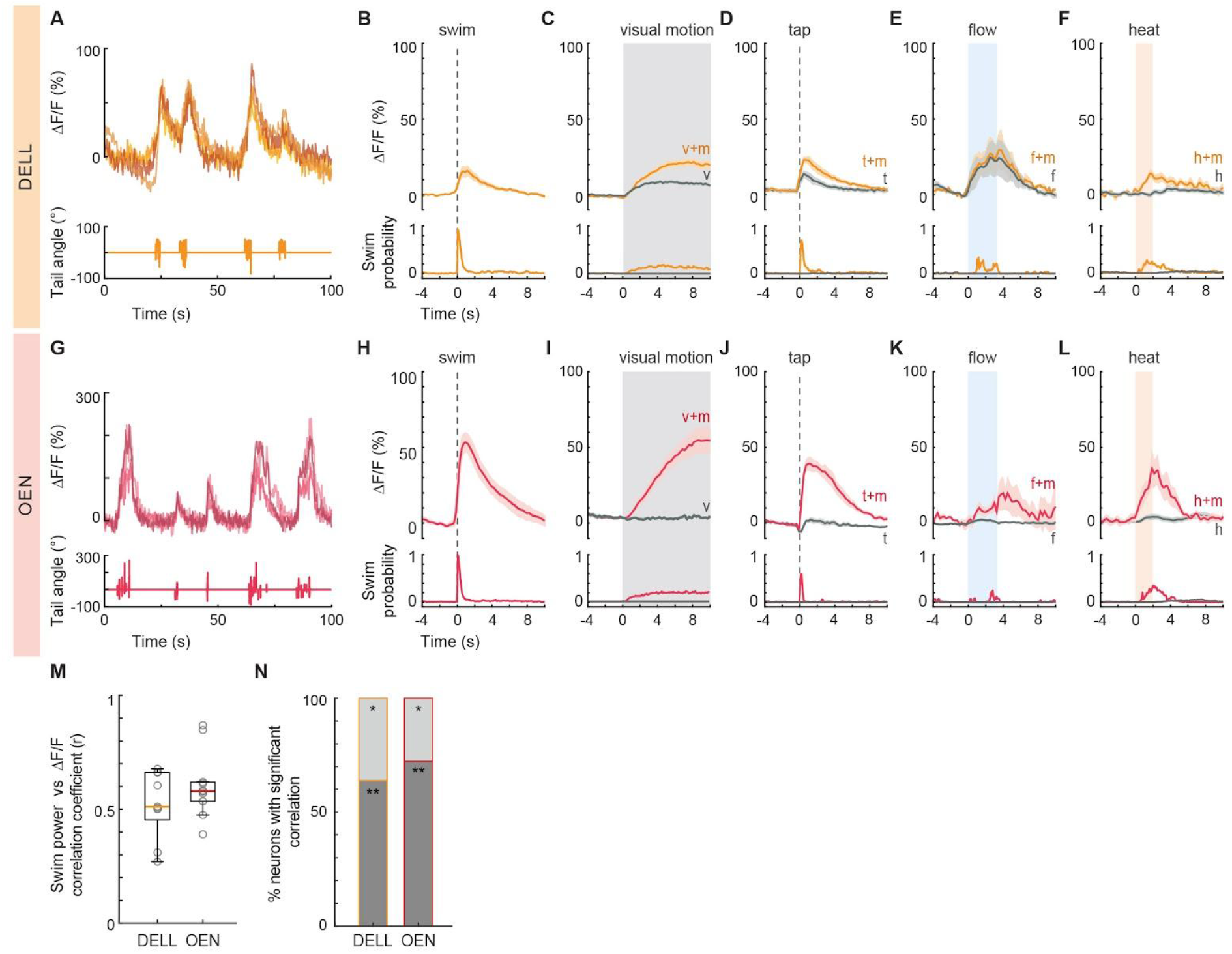
Activity of efferent nuclei during locomotion and in response to diverse sensory stimuli. **(A)** Example traces of top: fluorescence activity (ΔF/F) of 4 DELL neurons in a single fish, and bottom: the concomitant cumulative tail curvature of the animal. **(B-F)** Top: Population activity (mean ΔF/F ± s.e.m.) of neurons in the DELL of fish expressing GCaMP6s. Bottom: swim probability. **(B)** Average swim-triggered responses (n=9 fish). **(C)** Average stimulus-triggered responses during periods of moving visual gratings that elicited (v+m, orange) and failed to elicit (v, gray) motor responses. (n= 19 fish expressing GCaMP6f and 6s). Pooled data from fish from Figures 3B and S2A. **(D)** Average stimulus-triggered responses to taps that elicited motor responses (t+m, orange) and taps that did not (t, gray) (n= 7 fish). **(E)** Average stimulus-triggered responses to water flow stimuli over the animal’s body that elicited motor responses (f+m, orange) and those that did not (f, gray) (n= 3 fish). While the flow delivery period lasted 3.3 s, it is possible that a residual stimulus persisted after flow delivery was terminated. **(F)** Average stimulus-triggered responses during laser-mediated heat delivery events that elicited motor responses (h+m, orange) and events that did not (h, gray) (n=10 fish). **(G)** Example traces of top: fluorescence activity (ΔF/F) of 4 OEN neurons in a single fish, and bottom: the concomitant tail curvature of the animal. **(H-L)** Top: Population activity (mean ΔF/F ± s.e.m.) of neurons in the OEN of fish expressing GCaMP6s. Bottom: swim probability. **(H)** Average swim-triggered responses (n=8 fish). **(I)** Average stimulus-triggered responses during periods of moving visual gratings that elicited (v+m, red) and failed to elicit (v, gray) motor responses. (n= 16 fish expressing GCaMP6f and 6s). Pooled data from fish from Figures 3H and S2H. **(J)** Average stimulus-triggered responses to taps that elicited motor responses (t+m, red) and taps that did not (t, gray) (n= 5 fish). **(K)** Average stimulus-triggered responses during flow stimuli that elicited motor responses (f+m, red) and those that did not (f, gray) (n= 5 fish). **(L)** Average stimulus-triggered responses during laser-mediated heat delivery events that elicited motor responses (h+m, gredrey) and events that did not (h, gray) (n=8 fish). **(M)** Box plots showing the mean Pearson’s coefficients correlating the power of single swim bouts with the concurrent neuronal activity of DELL or OEN populations in individual fish (circles, n= 9 and n=10 fish expressing GCaMP6f, respectively). Medians shown in color. Swim power was defined as the integral of the absolute tail curvature trace for single bouts, and neuronal activity as the integral of the calcium transient during each corresponding bout. Plots derived from single-neuron analysis shown in Figures S2G and S2P. **(N)** Percentage of cells per fish whose activity was significantly correlated with swim power. (DELL: n=9, OEN: n=10 fish; ** dark gray p < 0.001, * light gray p < 0.05).

We next focused our attention on the OEN and observed that these cholinergic neurons also exhibit elevated activity during swims and startle responses (Figures 3G-H, 3J and S2H-I). However, in stark contrast to the DELL, OEN neurons responded exclusively during motor events and were not activated by any sensory stimulus in the absence of locomotion (Figures 3I-L and S2I). In accordance with this observation, ablation of the lateral line by copper treatment did not affect OEN responses in any way (Figure S2J), and OEN activity was indistinguishable during spontaneous and visually-evoked swims (Figure S2K). This pronounced coincidence with—and selectivity for—motor activity provides support for a primary role of this nucleus in efference copy transmission.

Since different behaviors generate different patterns of reafferent stimulation, and would thus require different patterns of efferent cancellation, we tested whether the activity of the OEN subnuclei differs when the animal executes distinct motor programs. To that end, we compared activity evoked by startles and swim bouts across the three OEN subnuclei. These two behaviors differ in that short-latency (<40 ms) startle responses are generated by Mauthner array cells (Foreman and Eaton, 1993; Liu and Fetcho, 1999; O’Malley et al., 1996), while routine swims are controlled by other reticulospinal neurons (Orger et al., 2008). We found that in spite of this distinct segregation of motor control units, OEN neurons in all subnuclei are synchronously active both during swims and startle responses (Figures S2L and S2M). This suggests that all three OEN subgroups can be thought of as one functional unit for the behaviors tested.

Finally, since accurate reafferent cancellation requires a matching of efferent and reafferent activity patterns, we examined whether increased motor vigor, which naturally elicits stronger reafferent activation, is correlated with stronger corollary neural activity in individual efferent neurons. Another possibility is that mechanosensation is completely silenced during locomotion, in which case efferent activity would serve as a simple gating signal and need not be adjusted to the strength of the motor output. We therefore calculated the correlation between swim strength and activity in individual neurons of both efferent nuclei, and asked specifically how well various kinematic features of the motor event match with the strength of efferent activity. Specifically, we correlated total swim power, as well as frequency and amplitude of tail beats with the integral of the fluorescent activity. All neurons in both nuclei were well-correlated with swim power (Figures 3M-N, S2G and S2P), while frequency and amplitude showed much weaker correlation coefficients (Figures S2E-F and S2N-O), suggesting that efferent activity scales with total vigor rather than with individual swim components. These results indicate that already at the zebrafish larval stage, neural mechanisms are in place that allow for a graded subtraction rather than a complete silencing of the reafferent sensory signals.

In summary, both DELL and OEN exhibit graded motor-correlated activity during swims and startle responses. DELL neurons, however, are also activated by visual and mechanosensory stimuli, while OEN neurons do not show any sensory-dependent excitation, making them the more likely mediators of efference copies to the lateral line.

### Ultrastructure

Our anatomical and functional results show that descending inputs from the brain reach the lateral line and influence sensory processing. To better understand the flow of information within this microcircuit, we set out to determine the precise connectivity patterns of all participating cells and neurites within a neuromast. We collected a complete serial section electron microscopy (ssEM) data volume of a neuromast and its surrounding tissue, and segmented the entire population of cells and neurites at this convergence zone (Movie S1). The animal used for this analysis expressed RFP in DELL neurons and GFP in OEN neurons, and was imaged using confocal microscopy prior to fixation for EM. These confocal image volumes allowed us to generate a detailed idiosyncratic anatomical footprint of each efferent neuron’s axonal branching pattern in and around the neuromast region. By comparing the fluorescent branching patterns with those of the segmented efferent neurites from the corresponding EM volume, we could unambiguously assign cell-type identities to the segmented structures in the EM data set (Movie S2).

In a single neuromast, we found 7 pairs of hair cells of opposing polarity (half rostro-caudal and half caudo-rostral). Since sterocilia in the hair cell bundle are organized in increasing length towards the kinocilium, polarity was determined by noting this asymmetry. We also detected multiple neurites: 6 afferent neurites selectively targeting one of the hair cell polarities, 3 axonal arbors originating from three separate DELL neurons and 2 axonal arbors originating from distinct OEN neurons (Figure 4A; Movie S3). Additionally, two neurites lacked counterparts in the confocal microscopy data, so their identities or origin could not be assigned (Movie S3). The complete matrix quantifying the pairwise connectivity of all circuit elements is shown in Figure 4G.

**Figure 4.**
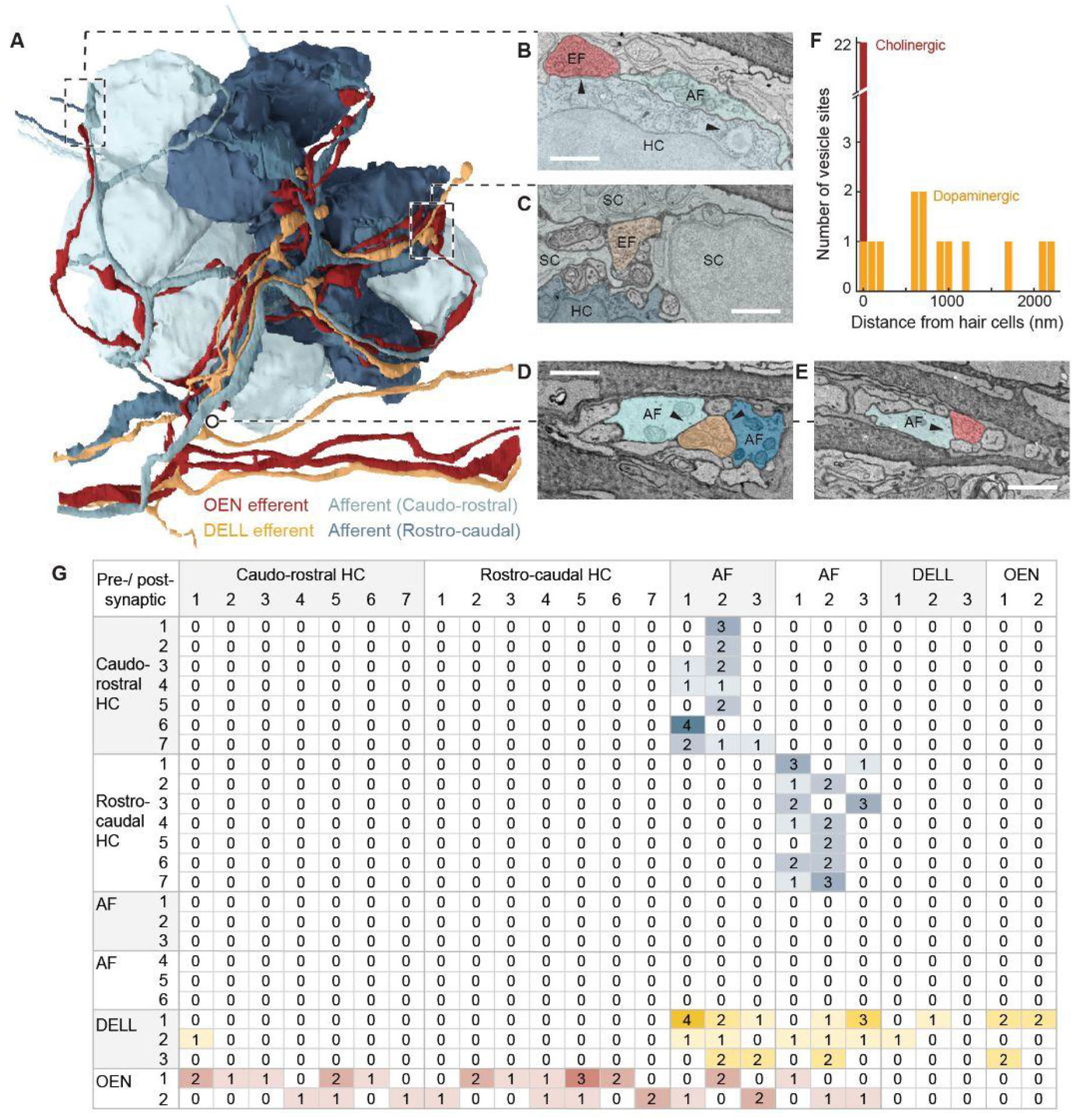
Volumetric reconstruction of a posterior lateral line neuromast in a 5 dpf fish from serial section electron microscopy data. **(A)** Innervation of a posterior lateral line neuromast. Neuron identities were assigned by correlating anatomical ssEM data to fluorescent labeling of efferent types in the same neuromast. This neuromast consists of 14 hair cells belonging to two equal populations of opposing hair-bundle polarities (rostro-caudal and caudo-rostral). **(B-E)** EM details showing examples of specific connections. **(B)** Hair cells (HC) and afferent neurons (AF) connect via ribbon synapses (right arrowhead). An efferent (EF) OEN terminal containing synaptic vesicles abuts the base of the hair cell, which contains postsynaptic membrane specializations (left arrowhead). **(C)** A vesicle-rich dopaminergic efferent terminal in the proximity of a hair cell. Note the distance between the efferent membrane and the hair cell, as well as the presence of intercalated support cells (SC). **(D)** A DELL efferent axon with vesicle-filled profile within the axon bundle innervating the neuromast. The vesicle-filled profile is in close apposition to afferent neurites carrying information from both polarities. **(E)** An OEN efferent axon with vesicle release sites, in close apposition to an afferent neurite within the axon bundle that innervates the neuromast. **(F)** Histogram showing the distances between DELL (orange) or OEN (red) vesicle sites and the closest hair cell. **(G)** Connectivity matrix summarizing the tally of all synaptic contacts observed in this neuromast (see methods for quantification).

The simplest hypothesis about the “micro-connectome” within this region is that hair cells form excitatory connections onto the afferent terminals of sensory neurons, whereas DELL and OEN terminals contact the hair cells with dopaminergic and cholinergic synapses respectively. Previous ultrastructural studies in zebrafish uncovered that hair cells connect with afferent sensory neurons via ribbon synapses (Bricaud et al., 2001; Dow et al., 2018; Metcalfe et al., 1985), and that afferent neurons selectively innervate hair cells of a single polarity (Faucherre et al., 2009; Nagiel et al., 2008). Our ultrastructural volumetric analysis corroborated this specificity: three of the afferent neurites received inputs from rostro-caudally-tuned hair cells exclusively, and the other three were targeted by hair cells of the opposite polarity (Figures 4A and 4G).

With respect to efferent innervation, we found that all hair cells receive at least one OEN input. OEN neurons indiscriminately innervate hair cells of both polarities, and partner choices seem to be stochastic: ten of the fourteen hair cells were targeted by one OEN neuron and seven by the other (Figure 4B, 4F and 4G). On the other hand, and in agreement with previous light-microscopy studies, vesicle-rich DELL terminals are more frequently found in the general vicinity, but not in close apposition to hair cells, often surrounded by support cells (Figure 4C and 4F; Movie S3) (Toro et al., 2015). We only identified a single close-range contact between a DELL axon and a hair cell (Figure 4F and 4G).

Extending our analysis to include the afferent neurites as potential targets of efferent innervation, we found that all of the fourteen vesicle-filled profiles from the three DELL axons abut an afferent neurite and they do so indiscriminately of their target’s polarity (Figures 4D and 4G). These profiles exist within the neuromast, and also further away, within the axon bundle (Figures 4A, 4C and 4D). This suggests that the dopaminergic synaptic sites are not necessarily diffuse, but can specifically modulate the sensitivity of the initial segment of the sensory dendrites. In contrast, OEN innervation of sensory afferents is less frequent, implying that the role of the OEN is largely dedicated to shaping hair cell sensitivity directly (Figures 4E and 4G). Interestingly, we detected multiple DELL vesicle-filled profiles abuting other DELL and OEN axons, which further strengthens the idea that dopaminergic modulation is acting systemically (Figures 4G ad S3).

Overall, by combining confocal microscopy with volumetric EM-based segmentation analysis, we were able to enrich ultrastructural data sets that provide nanoscale resolution with fluorescent cellular-identity labels. By determining the small set of connectivity rules in this anatomically complex structure, our approach uncovered a circuit of remarkable functional simplicity. Briefly, (1) hair cells receive mechanosensory inputs and transmit them to afferent neurons in a polarity-specific manner, (2) efferent neurons do not form polarity-specific connections, (3) OEN axons broadly innervate hair cells, and some afferent neurites, and (3) DELL vesicle-filled profiles directly abut afferent and some efferent neurites, while also terminating at a distance that allows for paracrine modulation of hair cell function.

### Circuit perturbations

#### Afferent neurons of the posterior lateral line ganglion acquire motor-evoked responses following ablation of OEN, but not DELL, inputs

To test the causal role of both DELL and OEN neurons in the modulation of sensory processing during locomotion, we measured the activity of lateral line sensory neurons in the PLL ganglion before and after targeted laser ablation of each individual efferent population. If any of these nuclei indeed play an essential role in silencing reafferent activity during swims, then their selective removal should relieve the afferent pathway from efferent inhibition and uncover self-evoked activity during locomotion. In order to first quantitatively evaluate the potential consequences of these ablations, we estimated the total number of neurons that comprise each nucleus. We used a simple capture-recapture random sampling approach, which is a standard method to estimate the total number of individuals in a population (see methods). We found that the number of cells in each nucleus is surprisingly low: on average there are only three to four neurons on each side in the DELL, while the srOEN, rOEN, and cOEN contain only one, two and three neurons on each side respectively (Figures S4A and S4B). This small population size provides an ideal context for targeted ablation studies since it is easy to remove a significant fraction, if not all, of the neurons in the ensemble (Figure S4C).

To ensure the selective targeting of these neurons, we performed unilateral dye injections in the lateral line nerve, and allowed at least 48 hours for nerve regeneration (Pujol-Marti et al., 2014; Villegas et al., 2012). This procedure consistently labeled all three neural populations: DELL, OEN and PLLg neurons, and the distinctive anatomical locations of their somas allowed for straightforward targeting of the ablations. Further, since efferent axons do not cross the midline, we could image PLLg neurons on the contralateral side to the ablation as controls. Removal of DELL neurons did not significantly change PLLg activity during locomotion, indicating that this efferent nucleus does not play a role in the selective suppression of reafferent inputs (Figures 5A-C and S4D). However, response patterns in PLLg neurons displayed a slightly enhanced variability after DELL ablations on the ipsilateral side, as indicated by a qualitative increase in the variance across fish and neurons (Figures 5C and S4D). This suggests that dopaminergic efferents might have some stabilizing modulatory influence, the precise nature of which still needs to be elucidated.

**Figure 5.**
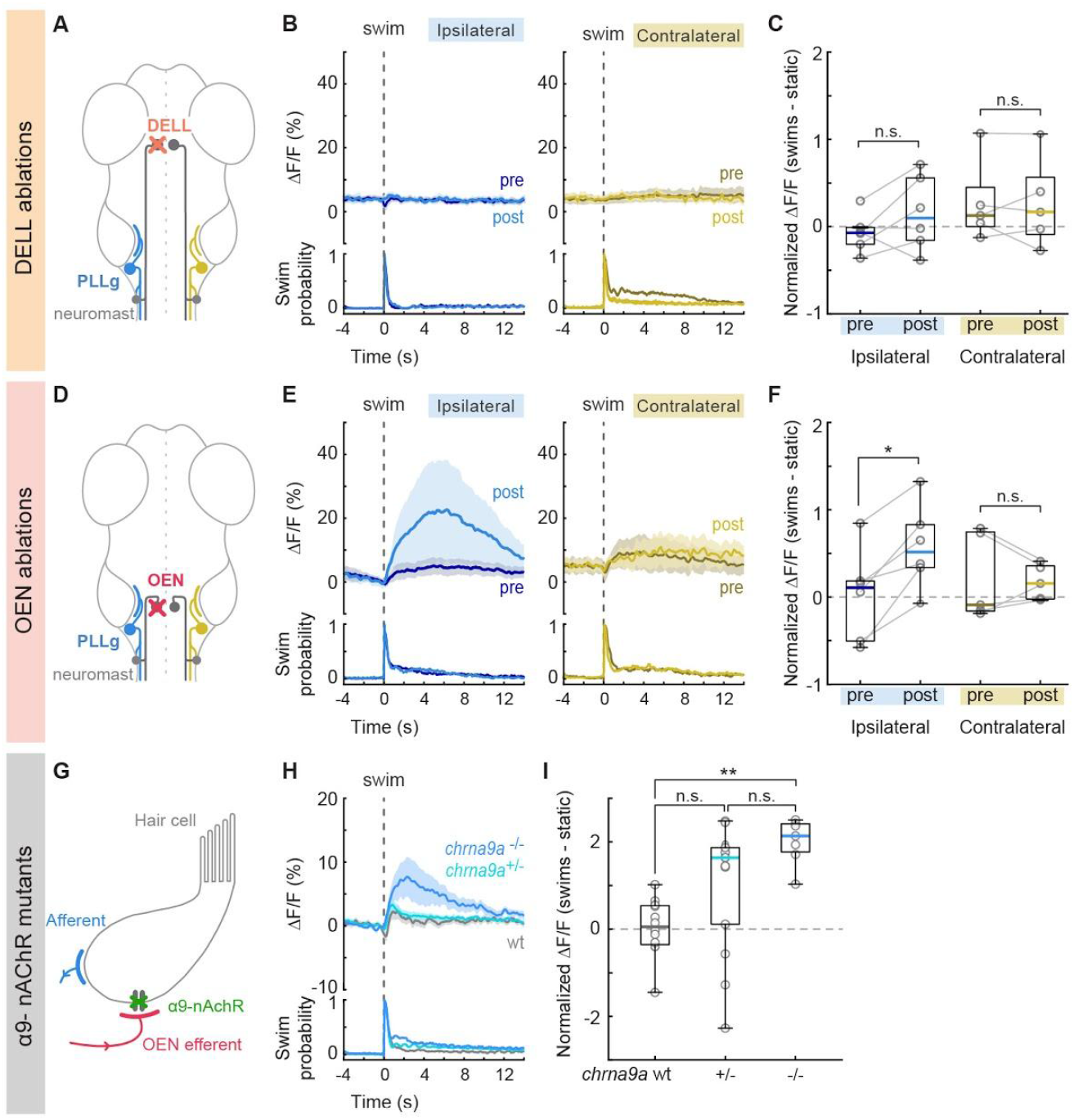
Swim-triggered activity of PLLg neurons after efferent nuclei ablation and in mutants lacking α9-nAChRs. **(A)** Circuit perturbation schematic. PLLg activity during swims was recorded before and after unilateral laser ablations of DELL neurons. Since DELL neurons do not cross the midline, contralateral PLL ganglia serve as controls. **(B)** Average swim-triggered responses (mean ΔF/F ± s.e.m.) before and after unilateral DELL ablations in neurons from PLLg ganglia in the ipsilateral (left, blue lines) or contralateral (right, yellow lines) side to the ablation. Bottom panels: swim probability. (n= fish) **(C)** Boxplots of z-scored population ΔF/F per fish showing the difference in neuronal activity during swimming and quiescent periods of PLL ganglia located ipsilaterally (blue) or contralaterally (yellow) to the ablation site. Medians shown in color. Differences were computed for each fish before and after DELL ablations. Population values correspond to the median of all PLLg neurons for a given fish (individual cell values can be found in Figure S4D). Paired 2-tailed Wilcoxon signed-rank test: p_ipsilateral_ = 0.2188, p_contralateral_= 1. **(D)** Circuit perturbation schematic. PLLg activity during swims was recorded before and after unilateral laser ablations of OEN neurons. OEN axons also do not cross the midline and therefore contralateral PLL ganglia serve as controls. **(E)** Average swim-triggered responses (mean ΔF/F ± s.e.m.) before and after unilateral OEN ablations in neurons from PLLg ganglia in the ipsilateral (left, blue lines) or contralateral (right, yellow lines) side to the ablation. Bottom panels: swim probability. (n= 6 and 5 fish, respectively). **(F)** Boxplots of z-scored population ΔF/F per fish showing the difference in neuronal activity during swimming and quiescent periods of PLL ganglia located ipsilaterally (blue) or contralaterally (yellow) to the ablation site. Medians shown in color. Differences were computed for each fish before and after OEN ablations. Population values correspond to the median of all PLLg neurons for a given fish (individual cell values can be found in Figure S3E). Paired 2-tailed Wilcoxon signed-rank test: p_ipsilateral_ = 0.0313, p_contralateral_= 0.8125. **(G)** Circuit perturbation schematic. Hair cells receive cholinergic inputs from OEN efferent neurons (red). To test whether α9-nAChRs mediate cholinergic inhibition, we generated *chrna9a* mutant zebrafish. If hair cell inhibition is compromised, this should be also observed in the activity of PLLg afferent neurons. **(H)** Average swim-triggered responses (mean ΔF/F ± s.e.m.) of PLLg neurons in *chrna9a*Δ*1049* homozygous (blue) and heterozygous (cyan) siblings, and wild-type controls (wt, gray), (n=8,14,12 fish). Wild-type fish correspond to one wt sibling and all animals from Figure 2F. Bottom: swim probability. **(I)** Boxplots of z-scored population ΔF/F per fish showing the difference in neuronal activity during swimming and quiescent periods in wildtype and mutant animals. Medians shown in color. (Kruskal–Wallis one-way analysis of variance, p= 0.0010. Unpaired 2-tailed Wilcoxon rank-sum test with post-hoc Bonferroni correction, p_wt:+/-_ =0.0252, p_+/-:-/-_ =0.0676, p_wt:-/-_ = 3.9692×10^−5^).

Next, we tested the effect of silencing the OEN efferent pathway on PLLg sensitivity. Here, we found that before ablations, sensory neurons in the labeled and control ganglia were silent both during periods of quiescence and during motor events, as expected (Figures 5D-F). Strikingly, after OEN ablations, a subset of neurons in the ipsilateral PLLg displayed robust responses during swimming, demonstrating that OEN input is necessary for sensory inhibition during locomotion (Figures 5E, 5F and S4E, blue). All neurons of the contralateral ganglion, on the other hand, continued to be silent during swim bouts (Figures 5E, 5F and S4E, yellow). The fact that a subset of ipsilateral PLLg neurons remained silent during swims indicates that some inhibition was preserved. This can be readily explained by the partial and stochastic nature of our ablation method, which on average left about three OEN neurons unlabeled and consequently intact in each animal. These functional results are fully in line with the ultrastructurally-determined connectivity patterns we have observed in Figure 4: individual hair cells do not receive inputs from all OEN neurons, but rather, are targeted stochastically by a small subset. Therefore, a partial removal of OEN neurons is expected to uncover activity only in a subset of PLLg neurons and leave the rest fully inhibited. Furthermore, this could also be explained by the observation that whilst all hair cells in a neuromast are mechanosensitive, a fraction of them are synaptically silent, and will fail to excite their postsynaptic afferent partners in the presence of a mechanosensory stimulus (Zhang et al., 2018).

#### Afferent neurons in the posterior lateral line ganglion exhibit motor-evoked stimulation in chrna9a knockout animals

In the mammalian cochlea, the inhibitory effect of cholinergic efferents onto hair cells is relayed through nicotinic receptor channels composed of specialized α9/α10 subunits (Elgoyhen et al., 1994, 2001). The efferent suppression of cochlear responses is lost in α9 knockout mice (Vetter et al., 1999). In the hair cells of zebrafish neuromasts, transcripts of the α9, but not the α10 subunit, have been detected (Erickson and Nicolson, 2015). We therefore asked whether specific removal of this receptor subunit would unmask PLLg activity during swim bouts, similar to the effect seen after OEN ablations. To this end, we generated *chrna9a* knockout zebrafish using CRISPR-Cas9 genome editing (Figures S4F and S4G). In homozygous *chrna9a* mutant animals, significant PLLg activity was observed during swim bouts, that was reduced in heterozygous siblings and absent in wild-type controls (Figures 5H and 5I). We confirmed that the re-emergent activity in the mutant animals originated, as expected, from the hair cells lacking functional nicotinic receptors, since this activity disappeared upon neuromast ablation using copper sulfate (Figure S4H). Furthermore, this activity was comparable in size to that observed in OEN ablated fish (Figures 5E and 5F), which suggests that both these perturbations disrupt the same synaptic pathway.

In summary, there is convergent evidence from ablation studies and genetic manipulations, that the cholinergic efferent pathway plays a critical role in silencing reafferent activity in the lateral line, and that this inhibitory action is precisely synchronized with the occurrence of locomotor events. The role of the dopaminergic pathway, on the other hand, is less clear. We did not observe any obvious modulatory effects of dopaminergic efferents on sensory neuron activity, but our ultrastructural analysis revealed synaptic contacts between efferent dopaminergic axons and afferent sensory neurites. It is possible that the modulatory effect of dopamine occurs over longer time-spans that precludes analysis in our experimental settings.

## Discussion

Reafferent sensation in any animal is invariably the consequence of a motor action, which is initiated by a neuronal command somewhere in the complex circuitry of the brain. This motor command is readily repurposed into a corollary discharge, or efference copy, which can be used to cancel out the expected self-generated reafferent stimulation. Understanding the principles behind this mechanism requires investigating it at the structural, functional and molecular levels. Here, we followed a multi-level approach to dissect the compensatory mechanisms that cancel out self-generated stimulation in the lateral line of the larval zebrafish. This comprehensive line of inquiry has revealed (1) how efferent responses are tuned to motor commands, (2) how synaptic connectivity supports the cancellation of reafferent information, and (3) how specific molecular receptors mediate the required inhibition.

### The Efferent Nuclei

We have identified the OEN and DELL as carriers of corollary discharges, as they both fire in strict synchrony with motor actions such as swims and escapes. However, only the OEN is shown to cancel out the predicted reafferent stimulation through direct inhibition of the hair cells via specialized nicotinic receptors.

The exact role of the DELL neurons remains unclear, but their dopaminergic nature together with their targeting of afferent neurites is suggestive of a role in sensory processing modulation. Furthermore, previous work has shown that dopamine can have a sensitizing effect on neuromast hair cells via the D1b receptor (Toro et al., 2015). The authors, however, did not detect D1b receptor expression in any of the neurons that innervate hair cells, so the molecular players in the synapses between DELL axons and afferent neurites are still unknown. Since DELL neurons exhibit increased activity during locomotion, and also in response to sensory stimuli such as moving visual gratings, flow and waves elicited by taps (Figures 3A-F), their effects are dependent on both motor-state and salient sensory stimulation (Reinig et al., 2017). By virtue of its action through G-protein coupled receptors, the effects of dopamine likely outlast the brief occurrences of swims or stimuli—leaving the sensory system in a more receptive state during the inter-bout periods or following stimuli that might require subsequent behavioral responses. Additionally, DELL neurons send collaterals to the spinal cord, and have been shown to affect bout frequency, indicating a role in the regulation of spinal circuit excitability (Barrios et al., 2020; Jay et al., 2015). Taken together, these observations suggest that DELL neurons serve a dual function in the control of basal threshold levels of both sensory and motor networks.

OEN neurons, on the other hand, transmit inhibitory efference copy signals directly onto hair cells to cancel out reafferent stimuli during locomotion (Lunsford et al., 2019; Pichler and Lagnado, 2020). Without such inhibition, hair cells habituate readily, which can render the animal intensitive to exafferent cues from the environment (Skandalis et al., 2021). Interestingly, individual OEN neurons broadcast their inhibitory signals to the lateral line and the inner ear, such that this small efferent population is used ubiquitously to cancel reafferent information in all sensory modalities that depend on hair cell function (Figure S1D and S1E). This simple strategy prevents informational ambiguity and most likely serves to suppress maladaptive triggering of behavioral responses.

### Subtraction vs Silencing

Mechanistically, reafferent cancellation could be achieved by a precise subtraction of the predicted reafferent signals during locomotion, or alternatively, by a complete shunting of hair cell excitability. The former, requires a finely-tuned and graduated inhibition that is matched to the strength of the reafferent signal, which in turn depends on the strength of the motor command. An exact subtraction of the reafferent signal, whose precision would need to be homeostatically adjusted to account for changes in the body of the animal (von Uckermann et al., 2016) or the properties of the environment (Bell, 1981; Bodznick et al., 1999; Kim et al., 2015), would allow for the detection of concomitant external cues, a necessity when animals move continuously and not in intermittent bouts. Since zebrafish transition from bout swimming to continuous locomotion during later stages in development, mechanisms might already be in place in the young larvae that favour subtraction rather than complete suppression. Our findings argue that the fundamental properties for subtraction are already in place. First, we observe that sensory neurons in the PLLg respond to exafferent flow stimulation during a swim, and that therefore the animal is not “blind” during locomotion (Figure 2B). Second, we show that OEN activity is highly correlated with the strength of motor output, consistent with a graded inhibition of the corresponding reafferent stimulation (Figures 3M 3N and S2P). Additionally, it has been shown that efferent inhibition of the neuromast in larval zebrafish is stronger in hair cells that are activated by forward motion, and that hair cells tuned to other perturbations are less affected (Pichler and Lagnado, 2020). Together, this suggests a selective tailoring of the inhibition to suppress motion-specific activation, a feature that will become necessary when the animal matures, starts swimming continuously and the nature of the reafferent stimulus becomes complex and long lasting.

### The Connectome of the Neuromast

Seminal work on the lateral line of zebrafish larvae provided the first EM images of a neuromast and established the existence of synaptic connections between hair cells and sensory afferents, and efferents and hair cells (Bricaud et al., 2001; Metcalfe et al., 1985). Subsequently, a comprehensive ultrastructural analysis of larval zebrafish neuromasts performed at an early developmental stage (∼3 dpf) rendered a first sketch of this interconnected microcircuit (Dow et al., 2018). In line with these studies, we confirm the close apposition of afferent terminals to ribbon synapses, as well as the precise selectivity of individual afferent neurons to hair cells of one specific polarity. However, Dow et al. found that in most neuromasts, a single efferent neurite contacts all hair cells of both polarities. In contrast, we find that multiple neurites of different types target some, but not all, hair cells or afferent neurites (Figure 4G). This discrepancy can be easily reconciled by taking into account that the previous study was agnostic to the neurotransmitter identity of the efferent neurites, and that at 3 dpf, the animal might present less refined connectivity patterns, as is commonly observed in other developing neural structures such as the neuromuscular junction (Tapia et al., 2012).

Since cholinergic terminals form direct synapses onto hair cells, the average distance between their respective membranes was, as expected, less than 100 nm (Figure 4F). For the majority of dopaminergic release sites, on the other hand, this distance exceeded 500 nm. This suggested at first glance, and in line with previous light microscopy work (Toro et al., 2015), that dopamine is released from diffuse synapses into the extracellular space of this densely interconnected region. However, upon closer examination of the completely reconstructed ultrastructural volume, we find that individual dopaminergic neurites appear to specifically target afferent terminals of sensory neurons, and they do so, similar to the cholinergic efferents, in a fashion that is independent of the polarity of their targets. Moreover, we have also uncovered axo-axonic connections between DELL neurons and other DELL or OEN axons (Figure 4G and S3). The nature and function of these dopaminergic connections remain to be elucidated.

Based on this ultrastructural analysis, we conclude that both efferent nuclei target both polarities of hair cells or associated afferent neurites indiscriminately, and appear to provide a global modulatory signal that is independent of the direction of the mechanosensory input.

### α9-nAChR Gene Knockouts

Our experiments using knockout larvae establish that the α9 cholinergic receptor subunit, which is also conserved in the hair cells of the mammalian inner ear, is pivotal for the suppression of reafferent mechanosensation in the neuromast. Recent work has determined that α9 subunits are expressed in zebrafish neuromasts, and that these can form functional homomeric acetylcholine-sensitive receptors in heterologous preparations (Carpaneto Freixas et al., 2021). Furthermore, using calcium imaging, this study showed that application of acetylcholine reduces calcium signals evoked by hair cell stimulation, and that this effect is blocked by apamin, a selective blocker of small-conductance calcium-activated potassium (SK) channels. The emerging view is that reafferent inhibition relies on acetylcholine-mediated calcium entry through α9 receptors, which subsequently activates calcium-dependent SK channels to hyperpolarize the hair cells. Whether this mechanism is also true for the inhibition afferent neurites remains to be determined. These observations are also consistent with results from the lateral line of *Xenopus laevis (Dawkins et al*., *2005)* and underscores the usefulness of dissecting the function of the lateral line of aquatic organisms, validating it as a more accessible model system for further studies of vertebrate hair cell function, also in the context of prevalent human pathologies related to the inner ear.

### A Complete Model of Local Microcircuitry

We show here that larval zebrafish possess parallel descending inputs that can differentially influence mechanosensory processing at early stages within the sensory pathway. Cholinergic signals from the hindbrain transmit efference copy signals that cancel out self-generated stimulation during locomotion, while dopaminergic signals from the hypothalamus may have a more complex role in modulating sensitivity in a broader behavioral context. These results allow us to propose a complete model—spanning circuit connectivity, neuronal function and synaptic receptors—of the circuit underlying reafferent mechanosensation in larval zebrafish.

## Author contributions

I.O. and F.E. conceived of the project and designed the experiments with contributions from R.P., M.H. and P.O. I.O. performed the experiments, using software written by R.P. in the behavioral set-up, and in collaboration with M.H. for experiments with heat delivery, and M.N. with flow. M.D.P. obtained and analyzed the neuromast connectome with support from J.B-W. for tissue preparation and tracing, J.B-W. and R.S. for data acquisition, A.P. for image alignment and volume reconstruction, and J.L. for infrastructure and general supervision. J.A.G. generated the *chrna9a* mutants. I.O. analyzed the data, and wrote the paper with F.E. All authors read and commented on the manuscript.

## Acknowledgments

We thank J. Miller and K. Hurley for fish maintenance and care, Douglas Richardson and the Harvard Center for Biological Imaging for microscopy infrastructure and advice, Renate Hellmiss and Mariana Grünthal for support with graphical design. We are grateful to Isaac Bianco, Alex Schier, Nao Uchida, Aravi Samuel, and Chris Harvey for valuable and helpful discussions, Polina Kehayova for resource management, and to Eva Naumann and Julian Arni for critical reading of the manuscript. This research was supported by funding from the National Institutes of Health (U19NS104653, R43 OD024879, 2R44OD024879), the National Science Foundation (IIS-1912293), the Simons Foundation (SCGB 542973), and the Human Frontier Science Program (RGP0033/2014) awarded to FE. IO was supported by the Human Frontier Science Program (LT000805/2019-L) and the European Molecular Biology Organization (ALTF 202-2019). RP was supported by the Max Planck Foundation and by the Deutsche Forschungsgemeinschaft (DFG, German Research Foundation) under Germany’s Excellence Strategy within the framework of the Munich Cluster for Systems Neurology (EXC 2145 SyN-ergy – ID 390857198).

## Methods

### Zebrafish

Zebrafish (*Danio rerio*) between 4-9 days-post-fertilization (dpf) were used for all experiments. Transgenic lines used as anatomical markers include: Tg(Isl1:GFP) to visualize OEN neurons (Higashijima et al., 2000), ETvmat2:GFP and Tg(DAT:GFP) to label DELL neurons (Wen et al., 2008; Xi et al., 2011), Tg(Brn3c:GFP) to tag hair cells of the lateral line and inner ear (Xiao et al., 2005), Tg(HGn39D:GFP) to mark lateral line primary sensory neurons (Faucherre et al., 2009), and Tg(elavl3:H2B-mCherry) to label neuronal nuclei. For functional imaging experiments, larvae homozygous for the Tg(elavl3:GCaMP6s) or Tg(elavl3:GCaMP6f) were used. In most cases, the animals were also homozygous for the mitfa^w2/w2^ skin-pigmentation (Lister et al., 1999). Fish were raised in fish facility water on a 14/ 10 hr light/dark cycle at 28 °C. Larvae were fed Paramecia daily from 5 dpf. Animal handling and experimental procedures were approved by the Harvard University Standing Committee on the Use of Animals in Research and Training.

### Dye labeling of efferent neurons

Larvae were anesthetized with 0.005 % MS-222 (Sigma) and embedded in low melting-point agarose on their sides. Sharpened tungsten needles were used to lesion the lateral line nerve at the level of neuromast L1 and to deposit labeled dextran crystals (Texas Red, Alexa Fluor 647, or Cascade Blue, 3,000 MW, Invitrogen). Fish were unembedded and allowed to recover for at least 24hs before proceeding with further experimentation.

### Capture-recapture random sampling

Neurons were tagged using the dye labelling technique described above. The ‘capture’ step involved injecting Cascade Blue-conjugated dextrans (3,000 MW, Invitrogen) at the level of the L1 neuromast, at 5 dpf. 48 hs were allowed for nerve regeneration (Villegas et al., 2012). In the ‘recapture’ step, the injection was repeated in the same location with Alexa Fluor 647-conjugated dextrans (3,000 MW, Invitrogen). The number of neurons labeled with the blue, far red and with both fluorophores were tallied 24 hs later and used to calculate the total population size using Chapman’s estimator (Chapman, 1954):

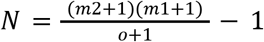, where N= total population size, m1= number of neurons marked the first time, m2= number of neurons marked the second time and o= number of neurons marked both times (overlap). Chapman’s estimator was favored over Lincoln-Petersen’s to account for small population sizes and allow for cases where no neurons were marked in both colors. This approach assumes that the population is closed: that no neurons are born or die between labeling sessions. It also assumes independence of sampling: that the probability of being ‘recaptured’ is not influenced by being ‘captured’ the first time. We performed these experiments on double transgenic animals Tg(elavl3:H2B-mCherry) expressing pan-neuronal, nuclear-targeted mCherry to aid in counting neurons, and Tg(Isl1:GFP) to have anatomical landmarks to distinguish between the three hindbrain nuclei.

### Immunohistochemistry

24 hs hours following dye labeling of the lateral line, fish were fixed in 4 % paraformaldehyde (PFA) diluted in PBS containing 0.25 % Triton (PBT). They were then immunostained using standard procedures (Inoue and Wittbrodt, 2011). Briefly, fish were washed in PBT, incubated in 150 mM Tris-HCl, pH 9, for 15 min at 70 °C, washed in PBT, permeabilized in 1 % Proteinase-K during 30 min, washed in PBT, blocked in blocking solution of PBT containing, 2 % normal donkey serum (NDS), 1 % bovine serum albumin (BSA), 1 % dimethyl sulfoxide (DMSO) and then incubated overnight at 4°C in primary antibodies diluted in blocking solution (goat anti-ChAT, 1:200, Abcam). Fish were then washed in PBT, blocked for 1 hr and incubated overnight at 4 °C in secondary antibodies conjugated to Alexa fluorophores (donkey anti-goat, 1:1000, Abcam).

### Confocal imaging

Imaging of live or fixed tissue after electroporation or immunohistochemistry, respectively, was performed with an upright confocal microscope (Zeiss LSM780) containing a 20x/1.0-NA water-dipping objective.

### Focal electroporations

Focal electroporations were performed in double transgenic larvae encoding GFP in hair cells Tg(Brn3c:GFP), and in efferent nuclei by means of Tg(Isl1:GFP) or ETvmat2:GFP to target the OEN or DELL, respectively. Electroporations were performed as described in (Tawk et al., 2009). Briefly, larvae were anesthetized and embedded in low melting-point agarose dorsal-side up. Micropipettes with tip diameters of 1-2 μm were filled with a 1 μl solution of plasmid DNA in distilled water. pCS2 expression vectors encoding mCherry or tdTomato fused to the N-terminal motif of the Src-family kinase Lyn were used. Guided by GFP expression in landmark areas, micropipettes were placed near the efferent nuclei, which were visualized under epi-fluorescence on a compound microscope. A Grass stimulator (SD9, Grass Technologies) was used to deliver plasmids by means of 1-2 200 Hz voltage pulse trains lasting 250 ms in 1 s intervals. Pulses were 20 V in amplitude and 2 ms in duration. Following electroporation, larvae were allowed to recover overnight in fish facility water. The next day, larvae were screened and those containing labeled neurons were anesthetized and mounted in agarose to be imaged using a confocal microscope (Zeiss LSM780).

### In vivo 2-photon functional imaging

A custom built 2-photon microscope was used for all functional experiments. The laser, a Ti:Sapphire ultra-fast laser (MaiTai, Spectra-Physics) was tuned to 950 nm, and operated at an average laser power of 5-10 mW at sample. Images were collected by scanning frames at 4 Hz and consecutive planes were separated by 2 μm. Image acquisition was controlled using custom software written in LabView (National Instruments).

### Behavioral tracking during functional imaging

Larvae were embedded in 2 % low melting-point agarose in a 35 mm Petri dish one day before imaging. Once the agarose solidified, filtered fish facility water was added and the agarose around the tail was removed with a scalpel to allow for free movement of the tail. To image, the dish was placed on the microscope’s transparent stage, which was covered with a diffusive screen onto which visual stimuli were projected. The screen also had a small hole to make the fish’s tail visible and recordable from below. Animals were illuminated with infrared light-emitting diodes (wavelength: 850 nm) and recorded at 200 frames per second (or 100 for heat experiments) using an infrared-sensitive camera (Pike F032B, Allied Vision Technologies). The cumulative angle of the tail was computed online and recorded as in Portugues and Engert, 2011 to determine the start and ends of individual swim bouts. This was used to update visual feedback in experiments with optomotor gratings and for analysis of locomotion throughout this study. Acquisition and stimulus presentation were controlled by custom programs written in LabView (National Instruments).

### Stimulus delivery during functional imaging

Optomotor gratings: visual stimuli were projected onto the screen under the fish at 60 frames per second using a 3M MPro110 micro-projector fitted with a red long-pass filter (Kodak Wratten Nr. 25) to enable imaging and visual stimulation simultaneously (Portugues and Engert, 2011). The stimulus consisted of a square wave grating with a spatial period of 10 mm and 100 % contrast (darkest and lightest pixels possible). Per imaging plane, the grating alternated 6 times every 20 s between being static or moving at 10 mm/s in the caudo-rostral direction. If a swim bout was detected, the grating speed was adjusted online to deliver the appropriate visual feedback in a closed-loop fashion.

#### Flow

A custom-built syringe pump system was used to deliver filtered fish facility water through a zero-dead-volume perfusion pencil (AutoMate Scientific), which was placed near the head of the fish as in Wee et al., 2020. After 4 s of baseline, 500 µl of fish facility water was delivered for 3.3 s. This was repeated every 20 s, 4 times per imaging plane.

#### Taps

A solenoid was fixed to the microscope stage and triggered by voltage pulses (Lacoste et al., 2015). The strength of the pulse was empirically determined for each fish at the beginning of an experiment to ensure that the animals would not respond to every tap. 4 taps were delivered per imagining plane, with interstimulus intervals of 10 or 15 s to avoid habituation.

#### Heat

Heat stimuli were delivered as in Haesemeyer et al., 2018. A 1 W 980 nm fiber-coupled diode laser (Roithner, Austria) coupled into a collimator (Aistana Inc., USA) was placed under the objective pointing downward at the fish at an angle of 16.5°, at a distance of 4 mm in front and 1.2 mm above. Laser power was programmatically controlled via a laser diode driver (Thorlabs, USA). Heat was delivered for 2 s in 20 s blocks (9 s off, 2 s on, 9 s off), 10 times in each imaging plane. The laser power at sample during the 2s pulse was 350 mW, heating the larva to a temperature ∼30 °C, which is aversive but below the noxious threshold.

### Data analysis

Data analysis was performed using scripts written in MATLAB (MathWorks).

Images were segmented manually to define ROIs corresponding to individual neurons using VAST (Berger et al., 2018). Segmentation was performed on an ‘anatomical stack’ obtained by computing the mean of the time-series for each plane. OEN and DELL neurons were identified by lateral line dye labeling, and segmentation was therefore performed on anatomical stacks obtained from the Alexa Fluor 647 signal in the volume imaged during the functional recordings. The PLLg can be clearly identified in GCaMP expressing animals as a ganglion posterior to the ear, so afferent neurons were segmented from anatomical stacks arising from the calcium imaging data.

The time-series of the fluorescence signal F(t) for each neuron was computed as the mean intensity of all pixels comprising an ROI in each imaging frame. The proportional change in fluorescence (ΔF/F) at time t was calculated as: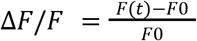, where F0 for any ROI is the 20th percentile of its entire fluorescence signal per plane. Perceptually uniform color maps developed by Ander Biguri were used to graph the average single-cell responses in Figure 2. *https://www.mathworks.com/matlabcentral/fileexchange/51986-perceptually-uniform-colormaps* To compute swim-triggered averages and for analyses correlating neuronal activity to different features of swim kinematics (Figure 3M-N, S2E-G and S2N-P), only ‘unitary’ bouts were taken into account. That is, bouts that occurred after at least 4 s from a previous bout and that were not followed by a subsequent bout within 2.5 s. The selected swim features were (1) Swim power, defined as the integral of the absolute tail curvature trace for an individual bouts, as in Ahrens et al., 2012. (2) Maximum amplitude, the maximum tail curvature achieved per bout and (3) Tail beat frequency, the inverse of the time between successive extreme tail positions in the same direction, as in Severi et al., 2014. Neuronal activity was defined as the integral of the calcium transient during each corresponding bout. Pearson’s coefficients correlating each bout feature with neuronal activity were calculated for each efferent neuron.

In order to quantify the change in PLLg activity after the ablation of efferent nuclei, and be able to compare effects across fish, the ΔF/F trace for each PLLg neuron was z-scored, and the average neuronal activity 4 s after a swim was subtracted to the average activity 4 s before a swim. A result of 0 indicates no difference in neuronal activity during swims or quiescent periods, whereas a positive result would indicate motor-correlated neuronal activity. Normalized differences were computed for each cell before and after ablations. Finally, the median of the normalized differences of all cells in a ganglion was calculated and used as the representative statistic for each animal (Figure 5C and 5F). A similar approach was taken when comparing PLLg activity in *chrna9a* mutants vs wildtypes (Figure 5). In this case, however, neurons of the PLLg were not individually segmented, but rather, the entire PLLg was considered a single ROI.

### Chemical ablation of neuromasts

Fish were incubated in 1 mM copper sulfate for 85 min and allowed to rest in fish facility water for 60 min. Only animals that showed complete neuromast ablation (assessed by DiASP staining: 0.5 mM in fish facility water for 15 min, as in Schuster and Ghysen, 2013 were used for additional functional imaging assays.

### Tissue preparation for ssSEM

A double transgenic zebrafish larva (Isl1:GFP and DAT:tdTomato) was used to visualize the two efferent nuclei with different fluorescent markers. At 5 dpf, the animal was anaesthetized in fish facility water containing 0.02 % (w/v) tricaine mesylate (MS-222, Sigma-Aldrich), and embedded in 2 % low melting-point agarose, lateral side up. The posterior lateral line neuromast (L1) was imaged under a confocal microscope to capture the pattern of innervation of the fluorescently-labeled efferent axons. L1 was selected because it lies above the swim bladder, which can be used as a landmark for subsequent steps. The animal was then prepared for fixation. Both eyes were enucleated in a dissection solution containing 0.02 % (w/v) tricaine mesylate (Ma et al., 2010), and the body was immediately transferred into cold fixative solution (2.5 % glutaraldehyde, 2 % paraformaldehyde, 3.5 % mannitol, 0.15 M sodium cacodylate buffer) and underwent two microwave fixation rounds (Tapia et al., 2012) followed by overnight incubation at 4 °C. Fixation continued for an additional 8 hrs with 2.5 % glutaraldehyde, 3.5 % mannitol, 0.15 M sodium cacodylate buffer and wash (0.15 M sodium cacodylate, 3×10 min). The sample was reduced with 0.8 %(w/v) sodium hydrosulfite in 60 % (v/v) 0.1 M sodium bicarbonate 40 % (v/v) 0.1 sodium carbonate buffer with 3 mM CaCl_2_ for 20 min and washed with buffer (3×10 min). Heavy metal staining was achieved as follows: 2 % osmium, 0.15 M sodium cacodylate (4 hrs RT, 4 °C overnight), 2.5 % (w/v) potassium ferrocyanide in 0.15 M sodium cacodylate (4 hrs RT, 4 °C overnight), wash with ddH_2_O water (3×10 min), filtered 1 % (w/v) thiocarbohydrazide (TCH) in ddH_2_O (1 hr, RT), wash with ddH2O and overnight incubation with 1 % uranyl acetate (UA) at 4 °C. The animal was then freed from the agarose and dehydrated with serial ethanol dilutions (25 %, 50 %, 75 %, 90 %, 100 %, 100 % ethanol in water, 10 mins each) and 100% propylene oxide (PPO, 2×10 min). The sample was infiltrated with a series dilution of LX-112 resin and PPO (25 %, 50 %, 75 % for 6 hrs and 100 % overnight), embedded at 60 °C for 72hrs. The cured block was trimmed, mounted, cut in 30 nm thick sections and imaged as in Hildebrand et al., 2017.

### Volumetric reconstructions from ssEM data

ssEM images were aligned using non-affine alignment through the FijiBento package on the Odyssey cluster supported by the FAS Division of Science, Research Computing Group at Harvard University (Saalfeld et al., 2012). Image segmentation was carried out manually using a custom volume annotation and segmentation tool. Segmented images were processed for 3D modeling with MATLAB and 3Ds Max for rendering.

VAST: https://software.rc.fas.harvard.edu/lichtman/vast.

### Laser ablations of efferent nuclei

Fish were subjected to unilateral dye injections in the lateral line at 4 dpf as described above, and allowed to recover in freely-swimming conditions for 48 hs to allow for nerve regeneration after injury (Pujol-Marti et al., 2014). At 6 dpf, fish were embedded in low-point melting agarose, and the agarose surrounding their tails was removed. Functional experiments were performed the following day. After a round of baseline functional experiments, we performed the ablation procedure as described in (Orger et al., 2008), with the exception that anesthesia was not used. Individual efferent neurons were targeted systematically by receiving 1-3 850 nm laser pulses of 1 ms, at 80 % laser power. Fish were then immediately used for functional experiments to test for the effects of ablations on sensory processing. On average, 4 OEN efferent neurons were ablated from the 8 that are hypothesized to exist (Figure S4B).

### Generation of *chrna9a* mutants using CRISPR–Cas9

Six Cas9 target sites within the *chrna9a* open reading frame were chosen using CHOPCHOP v2 (Labun et al., 2016) to generate single-guide RNAs (sgRNAs) for mutagenesis. DNA templates for transcription of sgRNAs were generated using oligo annealing and fill-in as previously described (Gagnon et al., 2014). Cas9 protein (NEB) was mixed with all six sgRNAs and injected into embryos at the 1-cell stage. Clutches from outcrosses of these injected fish were screened by PCR using flanking primers and sequencing was used to identify a 1049 bp deletion allele that deleted much of the third and fourth exons and generated a frameshift (Figure S4F). Founders were outcrossed repeatedly with Tg(elavl3:GCaMP6s) animals to reduce the likelihood of unlinked, off-target mutations affecting the *chrna9a* null genotype, and to be able to perform functional imaging experiments. Mutant fish were genotyped using a two primer PCR-based strategy. This mutant can be obtained from F. Engert (Harvard University). CRISPR target sites and primer sequences are listed below.

chrna9a_target1 GGACCCCCAGACACTAATGTGG

chrna9a_target2 AGAACTCTTGGTGATGGCAGGGG

chrna9a_target3 TGCATTCCTGACTATCAAAGGGG

chrna9a_target4 CGTACACAGTCCTGCTCAAGCGG

chrna9a_target5 GTGGAGCCAAGAAAGAGATGAGG

chrna9a_target6 GGGGAGAAGGTCTCGTTGGGGG

chrna9a_flanking_PCR_F GCTCAGTGCAGATGAGGAGG

chrna9a_flanking_PCR_R ACCATCAGCTGAAATACAGTCAGAG

## Statistics

All swim- and stimulus-triggered average plots show the mean ± s.e.m. across fish. Significance was determined using nonparametric statistical tests: the Wilcoxon signed-rank test for paired data, and the Wilcoxon rank-sum test for independent samples. Two-tailed p values are reported for all tests as follows: * p ≤ 0.05 and ** p ≤ 0.001. For correlation analysis, Pearson’s correlation was computed. When comparing fish populations of multiple genotypes (Figure 5I), the nonparametric Kruskal-Wallis one-way analysis of variance (ANOVA) tes was used, followed by Bonferroni corrected unpaired Wilcoxon rank-sum tests. In these cases, the genotype was determined post-hoc and disclosed after the functional data had been analyzed. No statistical methods were used to predetermine sample size.

**Movie S1. Annotated ssEM volume of a neuromast. Related to Figure 4**. Fly-through of the annotated ssEM stack of a posterior lateral line neuromast of a 5 dpf fish.

**Movie S2. Neuronal-type identification via correlation of confocal and EM volumes. Related to Figure 4**. Efferent neuronal identities were assigned by correlating EM images to confocal fluorescent images obtained prior to fixation. In addition to the assigned afferent and efferent neurons, there are two neurons with unassigned identities: (i) putative afferent neuron (blue) which does not form contacts of any kind, has no synaptic vesicles and becomes myelinated within the nerve bundle, and (ii) putative dopaminergic efferent neuron, which resides near the base of the hair cells, does not form contacts with the hair cells and has synaptic vesicles, but is not observed in the confocal image stack.

**Movie S3. Volumetric recontrunstruction of a neuromast. Related to Figure 4**. Full segmentation of the lateral line neuromast. All hair cells and innervating neurons have been densely segmented. Additionally, some support cells have been segmented (green shade) and the layer of skin collagen (blue) is shown.

---

**Figure S1.**
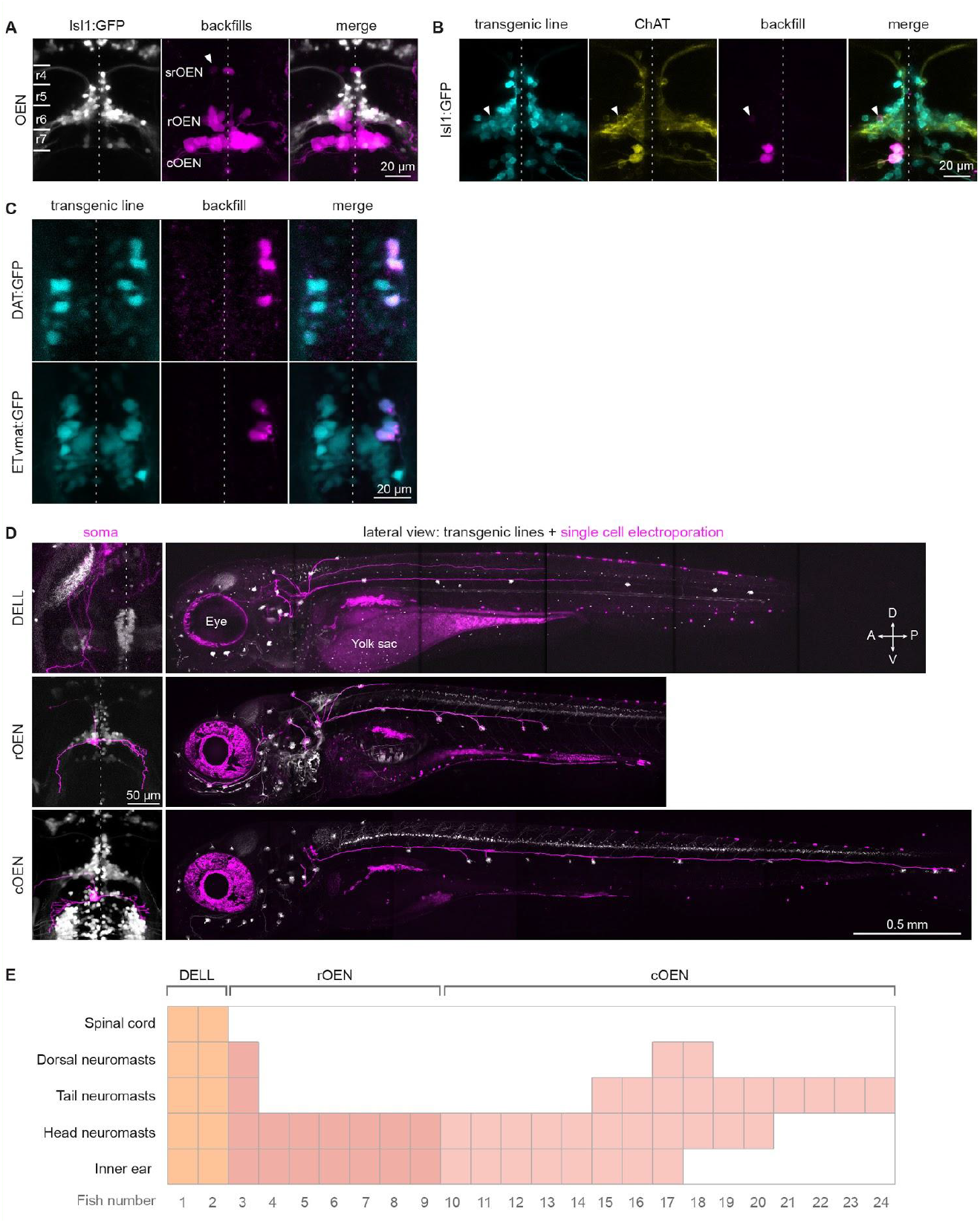
Characterization of neurotransmitter identity and target of efferent nuclei. Related to Figure 1. **(A)** The OEN is segregated into anatomically distinct subnuclei. Composite image showing all the neurons in the hindbrain labeled after dye injections of the lateral line nerve in Isl1:GFP larvae. Images were registered using anatomical landmarks made visible by GFP expression. The labeled neurons occupy three distinct positions along the antero-posterior axis spanning rhombomeres r4 to r7, and are named accordingly (caudal, rostral, and supra-rostral OEN). (n=17 fish, 6 bilateral). Dotted line: dorso-lateral midline. **(B)** Maximum intensity projections of confocal images of an Isl1:GFP larval hindbrain after a unilateral lateral line nerve dye injection and subsequent antibody stain against choline acetyltransferase (ChAT). Arrow: labeled neuron in the rOEN. **(C)** Maximum intensity projections of confocal images of the hypothalamus of transgenic larvae following unilateral lateral line nerve dye injections. Neurons labeled by the injections also express GFP under the Dopamine Active Transporter (DAT, top), and the Vesicular Monoamine Transporter (VMAT, bottom) promoters.**(D)** Focal electroporations of membrane-tagged fluorescent proteins reveal efferent neuron morphologies. Maximum intensity projections of confocal images showing the morphology of lateral line efferent neurons. Dorsal and lateral views of single cells (magenta) electroporated in double transgenic larvae expressing GFP in all hair cells, Tg(Brn3c:GFP), and in additional lines that label the efferent nuclei (gray). Top: DELL neurons in the ETvmat2:GFP background (5 dpf). Middle/bottom: rostral and caudal OEN neurons in Isl1:GFP fish (10 and 8 dpf, respectively). We were unable to successfully label a single srOEN neuron using this technique. Note: both the eyes and the yolk sac are auto-fluorescent and therefore also appear magenta. This signal is unrelated to the electroporations. **(E)** Matrix summarizing the observed target organs of individual efferent neurons belonging to different efferent nuclei and subnuclei.

**Figure S2.**
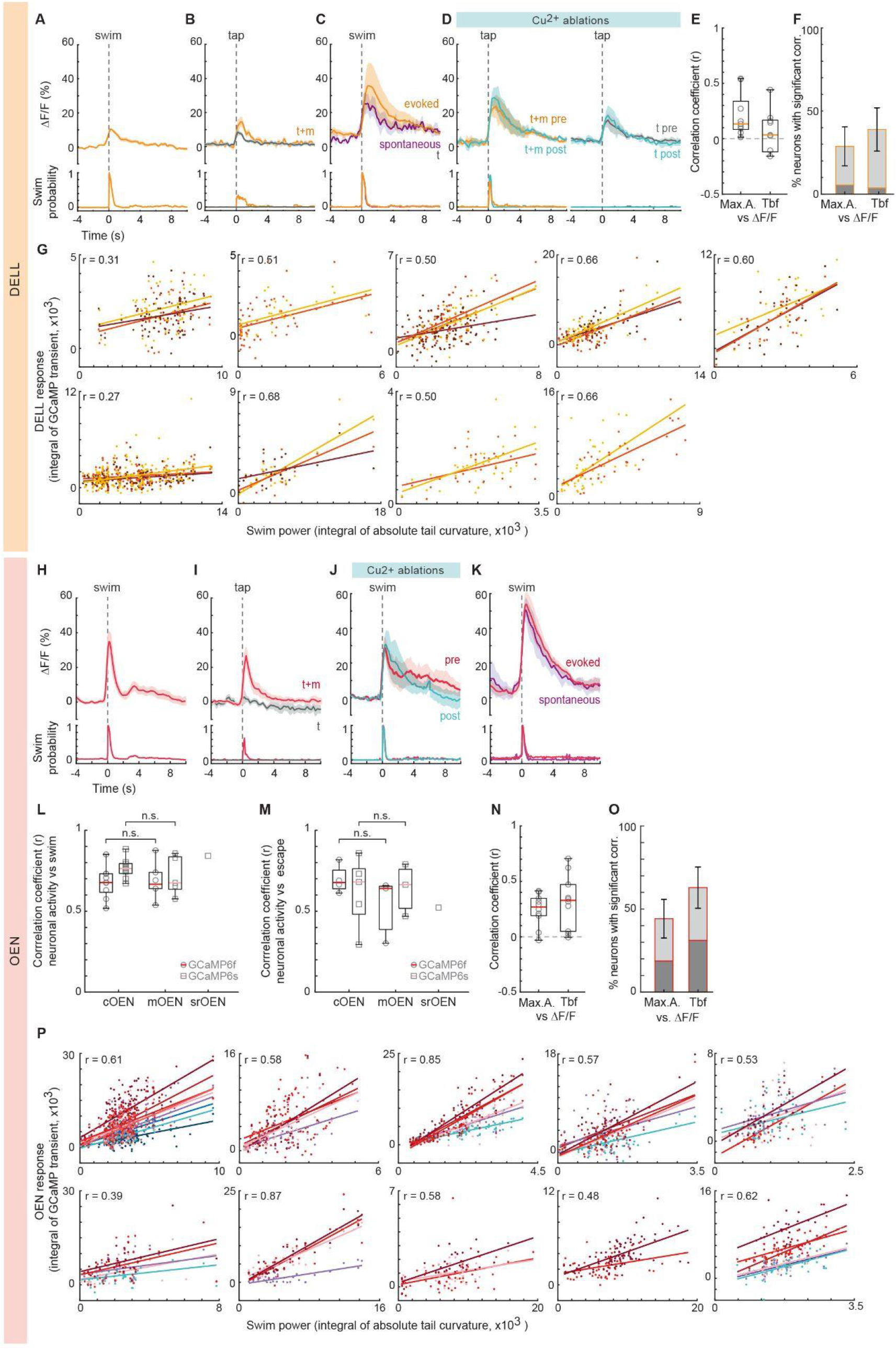
Activity of efferent nuclei during locomotion and in response to diverse sensory stimuli. Related to Figure 3. **(A-D)** Top: Population averages of DELL neuronal activity (mean ΔF/F ± s.e.m.). Bottom: swim probability. **(A)** Average swim-triggered neuronal responses (n=6 fish expressing GCaMP6f). **(B)** Average stimulus-triggered neuronal responses to taps that elicited motor responses (t+m, orange) and taps that did not (t, gray) (n= 5 fish expressing GCaMP6f). **(C)** Average swim-triggered neuronal responses after swims bouts that were spontaneous (purple) or evoked by a moving grating (orange) (n=5 fish expressing GCaMP6s). **(D)** Average stimulus-triggered neuronal responses during taps before and after ablation of the lateral line using copper sulfate. Responses are separated depending on whether the tap elicited (left) or did not elicit (right) motor responses (n= 3 fish expressing GCaMP6s/f). **(E)** Box plots showing the mean Pearson’s coefficients correlating maximum tail amplitude (Max. A.) or tail beat frequency (Tbf) of single bouts with the concurrent neuronal activity of DELL neurons in individual fish (gray circles, n= 9 fish expressing GCaMP6f). Median shown in color. **(F)** Mean percentage of DELL cells per fish whose activity was significantly correlated with maximum tail amplitude or tail beat frequency (n=9 fish, dark gray p < 0.001, light gray < 0.05, error bars: s.e.m). **(G)** Scatter plots showing the relationship between the power of each individual swim bout versus the intensity of the concurrent neuronal responses of all labeled DELL neurons in each of the 9 fish analyzed. Correlation coefficients and best-fit lines arising from linear regression were calculated for each neuron, and used to calculate the mean correlation coefficient (r) per fish. Swim power was defined as the integral of the absolute tail curvature trace for individual bouts, and neuronal activity as the integral of the calcium transient during each corresponding bout. **(H-K)** Top: Population averages of OEN neuronal activity (mean ΔF/F ± s.e.m.). Bottom: swim probability. **(H)** Average swim-triggered neuronal responses (n=9 fish expressing GCaMP6f). **(I)** Average stimulus-triggered neuronal responses to taps that elicited motor responses (t+m, red) and taps that did not (t, gray) (n= 5 fish expressing GCaMP6f). **(J)** Average swim-triggered neuronal responses before and after ablation of the lateral line by copper sulfate bath-application (n=6 fish expressing GCaMP6s/f). **(K)** Average swim-triggered neuronal responses after swims bouts that were spontaneous (purple) or evoked by a moving grating (red) (n=7 fish expressing GCaMP6s.) **(L)** Boxplots of mean Pearson’s coefficients relating swimming behavior and neuronal activity (ΔF/F) in different OEN subnuclei of fish expressing GCaMP6f (circles, c=9, r=6) or 6s (squares, c=8, r=5, sr=1). Medians shown in color. Locomotor behavior was convolved with a calcium kernel to account for calcium dynamics. (Pearson’s correlation, p<0.001; 2-tailed Wilcoxon rank-sum test, p= 0.86 and 0.52 for GcAMP6f and 6s, respectively.) **(M)** Boxplots of mean population Pearson’s coefficients relating short-latency escape behaviors and neuronal activity (ΔF/F) in different OEN subnuclei of fish expressing GCaMP6f (circles, c=5, m=3) or 6s (squares, c=5, r=3, sr=1). Median shown in color. Locomotor behavior was convolved with a calcium kernel to account for calcium dynamics.(Pearson’s correlation, p<0.001; 2-tailed Wilcoxon rank-sum test, p= 0.23 and 0.99 for GcAMP6f and 6s, respectively.) **(N)** Box plots of mean Pearson’s coefficients correlating maximum tail amplitude (Max.A.) or tail beat frequency (Tbf) of single bouts with the concurrent neuronal activity of OEN neurons of individual fish (gray squares, n= 10 fish expressing GCaMP6f). Median shown in color. **(O)** Mean percentage of OEN cells per fish whose activity was significantly correlated with maximum tail amplitude or tail beat frequency (n=10 fish, dark gray p < 0.001, light gray < 0.05, error bars: s.e.m). **(P)** Scatter plots showing the relationship between the power of all individual swim bouts versus the neuronal responses of all labeled OEN neurons in each of the 10 fish analyzed. Correlation coefficients and best-fit lines arising from linear regression were calculated for each neuron (shown in different shades), and used to calculate the mean correlation coefficient (r) per fish. Swim power was defined as the integral of the absolute tail curvature trace for individual bouts: a stationary tail has little curvature and thus power is ∼0, whereas an undulating tail has a positive absolute tail curvature, which increases as a function of motor strength. Neuronal activity was defined as the integral of the calcium transient during each corresponding bout.

**Figure S3.**
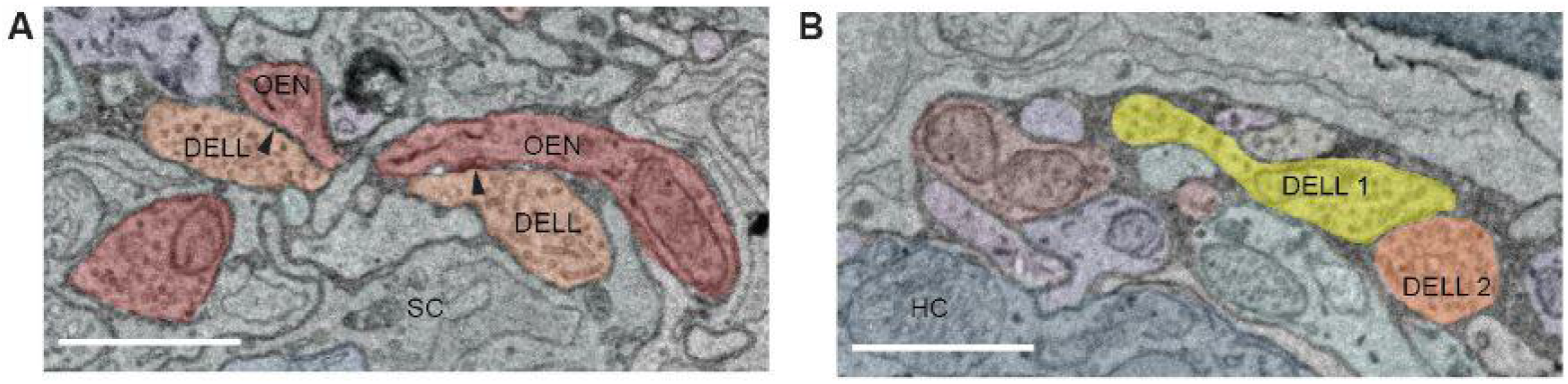
ssEM images of efferent to efferent connections in a posterior lateral line neuromast. Related to Figure 4. **(A)** DELL vesicle-filled profiles in close apposition to an OEN axon (arrowheads), surrounded by a support cell (SC). **(B)** Vesicle-filled profiles from two separate DELL neurons contact each other. A hair cell (HC) is also labeled for reference. Scale bars: 1 μm.

**Figure S4.**
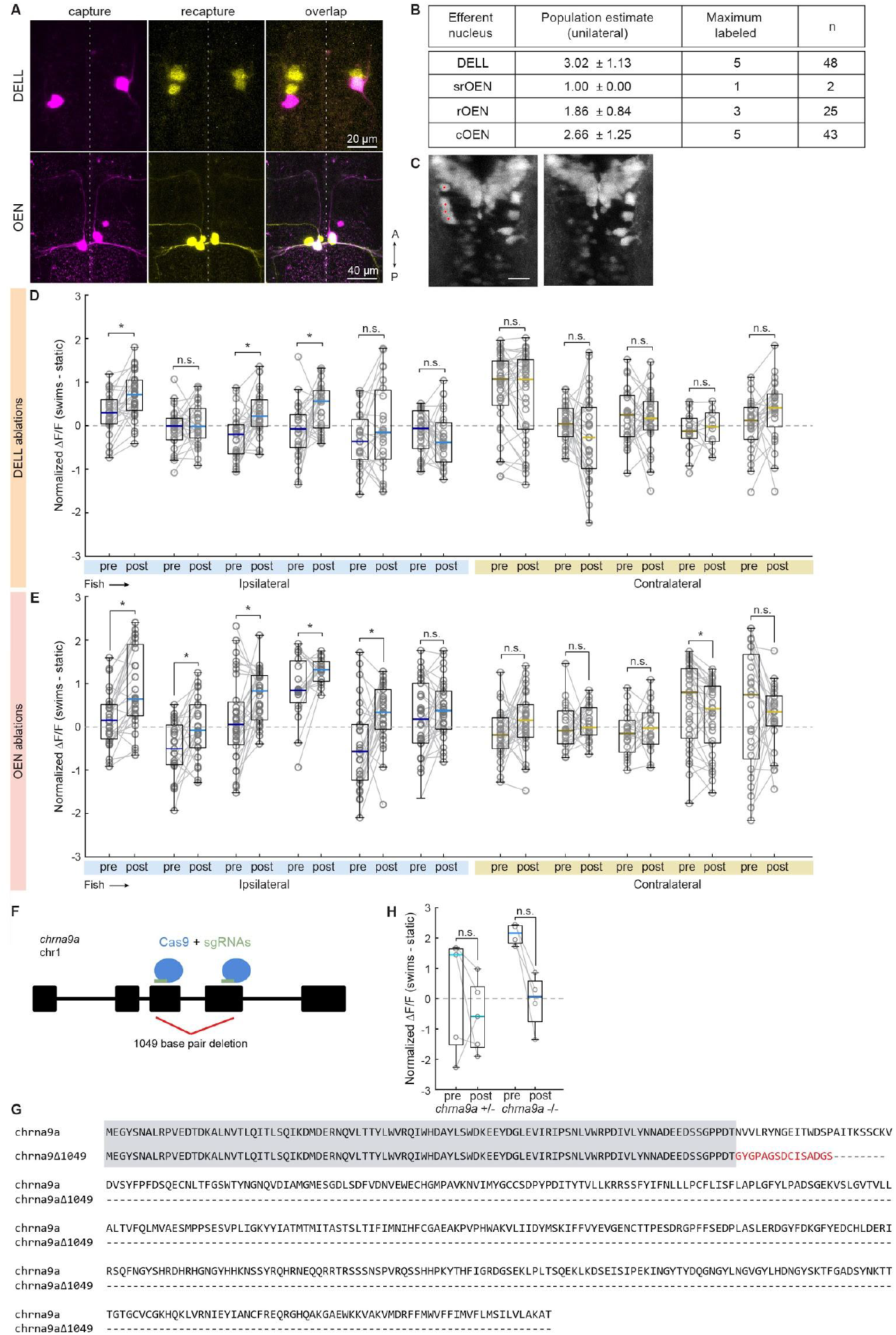
Efferent nuclei size estimation, effects of efferent nuclei laser ablation on PLLg activity and generation of mutants lacking α9-nAChRs. Related to Figure 5. **(A)** Dorsal projections of confocal images of fish that underwent consecutive dye injections of the lateral line nerve to estimate efferent population size. Top: DELL neurons, bottom: OEN neurons. **(B)** Estimated number of neurons comprising each efferent nucleus innervating the PLL. **(C)** Dorsal projections of confocal images of DELL neurons before and after laser ablations. Red dots indicate targeted neurons. **(D-E)** Boxplots of z-scored ΔF/F showing the difference in activity during swimming and quiescent periods of single neurons in PLL ganglia located ipsilaterally (blue) or contralaterally (yellow) to the ablation site. Medians shown in color. **(D)** Differences were computed for each neuron before and after DELL ablations. Paired 2-tailed Wilcoxon signed-rank test: p_ipsilateral_ = 0.3257,0.0714, 0.8854, 0.9219, 0.0814, p_contralateral_= 0.0014, 0.3673, 0.0014, 0.0107, 0.2238, 0.3158. **(E)** Differences were computed for each neuron before and after OEN ablations. Paired 2-tailed Wilcoxon signed-rank test: p_ipsilateral_ = 9.994×10^−4^, 0.0089, 0.0057, 0.0269, 0.0107, 0.1714, p_contralateral_= 0.0524, 0.6378, 0.3547, 0.0032, 0.5449. **(F)** Diagram of the *chrna9a* gene highlighting the CRISPR-Cas9 target sites in exons 3 and 4 and the 1049 base pair region deleted in the mutant. Introns depicted as thin lines. **(G)** Predicted amino acid sequence of the wild-type Chrna9a and mutant Chrna9aΔ1049 proteins. Shared sequences are highlighted in gray and amino acids changed due to frameshift in red.

